# Rapid, field-deployable nucleobase detection and identification using FnCas9

**DOI:** 10.1101/2020.04.07.028167

**Authors:** Mohd. Azhar, Rhythm Phutela, Asgar Hussain Ansari, Dipanjali Sinha, Namrata Sharma, Manoj Kumar, Meghali Aich, Saumya Sharma, Riya Rauthan, Khushboo Singhal, Harsha Lad, Pradeep Kumar Patra, Govind Makharia, Giriraj Ratan Chandak, Debojyoti Chakraborty, Souvik Maiti

## Abstract

Detection of pathogenic sequences or variants in DNA and RNA through a point-of-care diagnostic approach is valuable for rapid clinical prognosis. In recent times, CRISPR based detection of nucleic acids has provided an economical and quicker alternative to sequencing-based platforms which are often difficult to implement in the field. Here, we present FnCas9 Editor Linked Uniform Detection Assay (FELUDA) that employs a highly accurate enzymatic readout for detecting nucleotide sequences, identifying nucleobase identity and inferring zygosity with precision. We demonstrate that FELUDA output can be adapted to multiple signal detection platforms and can be quickly designed and deployed for versatile applications including rapid diagnosis during infectious disease outbreaks like COVID-19.

---

The rise of CRISPR Cas9 based approaches for biosensing nucleic acids has opened up a broad diagnostic portfolio for CRISPR products beyond their standard genome editing abilities^1,2^. In recent times, CRISPR components have been successfully used for detecting a wide variety of nucleic acid targets such as those obtained from pathogenic microorganisms or viruses and disease-causing mutations from various biological specimens^3-10^. At the heart of such a detection procedure lies the property of CRISPR proteins to accurately bind to target DNA or RNA, undergo conformational changes leading to cleavage of targets generating a reporter-based signal outcome^11-15^. To enable such a detection mechanism to be foolproof, sensitive and reproducible across a large variety of targets, the accuracy of DNA interrogation and subsequent enzyme activity is extremely critical, particularly when clinical decisions are to be made based on these results^1,2^.

Current technologies relying on using CRISPR components for nucleic acid detection can sense the identity of the target either through substrate cleavage mediated by an active CRISPR ribonucleoprotein (RNP) complex or by binding through a catalytically inactive RNP complex. Cleavage outcomes are then converted to a reporter-based readout with or without signal amplification. Among the CRISPR proteins that have been used so far, Cas9 and Cas12 or their inactive forms have been predominantly employed for detecting DNA sequences while Cas13 has been used for both DNA and RNA sequences. Each of these approaches has its own strengths and limitations that are related to sensitivity, specificity and read-out modes for an accurate diagnosis. The primary focus of these platforms is towards detection of low copy numbers of nucleic acids from body fluids of patients where signal amplification through collateral activity of fluorescent reporters has been proven to be advantageous. For genotyping individuals with high confidence, including careers of a single nucleotide variant (SNV), Cas12 requires individual sgRNA design and optimization for every variant and Cas13 requires conversion to an RNA template prior to detection, both of which increases the complexity and thus the time taken for diagnosis. Taken together, development of a detection pipeline based on a highly specific CRISPR protein with a direct binding or cleavage based readout can significantly increase the sensitivity of detection and reduce the time and cost of CRISPR based diagnostics (CRISPRDx), This is especially crucial for point-of-care (POC) applications where complex experimentation or setup of reaction components are not feasible.

We have recently reported a Cas9 ortholog from *Francisella novicida* (FnCas9) showing very high mismatch sensitivity both under *in vitro* and *in vivo* conditions^16-18^. This is based on its negligible binding affinity to substrates that harbor mismatches, a property that is distinct from engineered Cas proteins showing similar high specificity^19^. We reasoned that FnCas9 mediated DNA interrogation and subsequent cleavage can both be adapted for accurately identifying any SNVs provided that the fundamental mechanism of discrimination is consistent across all sequences (Figure 1A). We name this approach FnCas9 Editor Linked Uniform Detection Assay (FELUDA) and demonstrate its utility in various pathological conditions including genetic disorders and infectious diseases including disease outbreaks like COVID-19^20-22^.

**Fig. 1:**
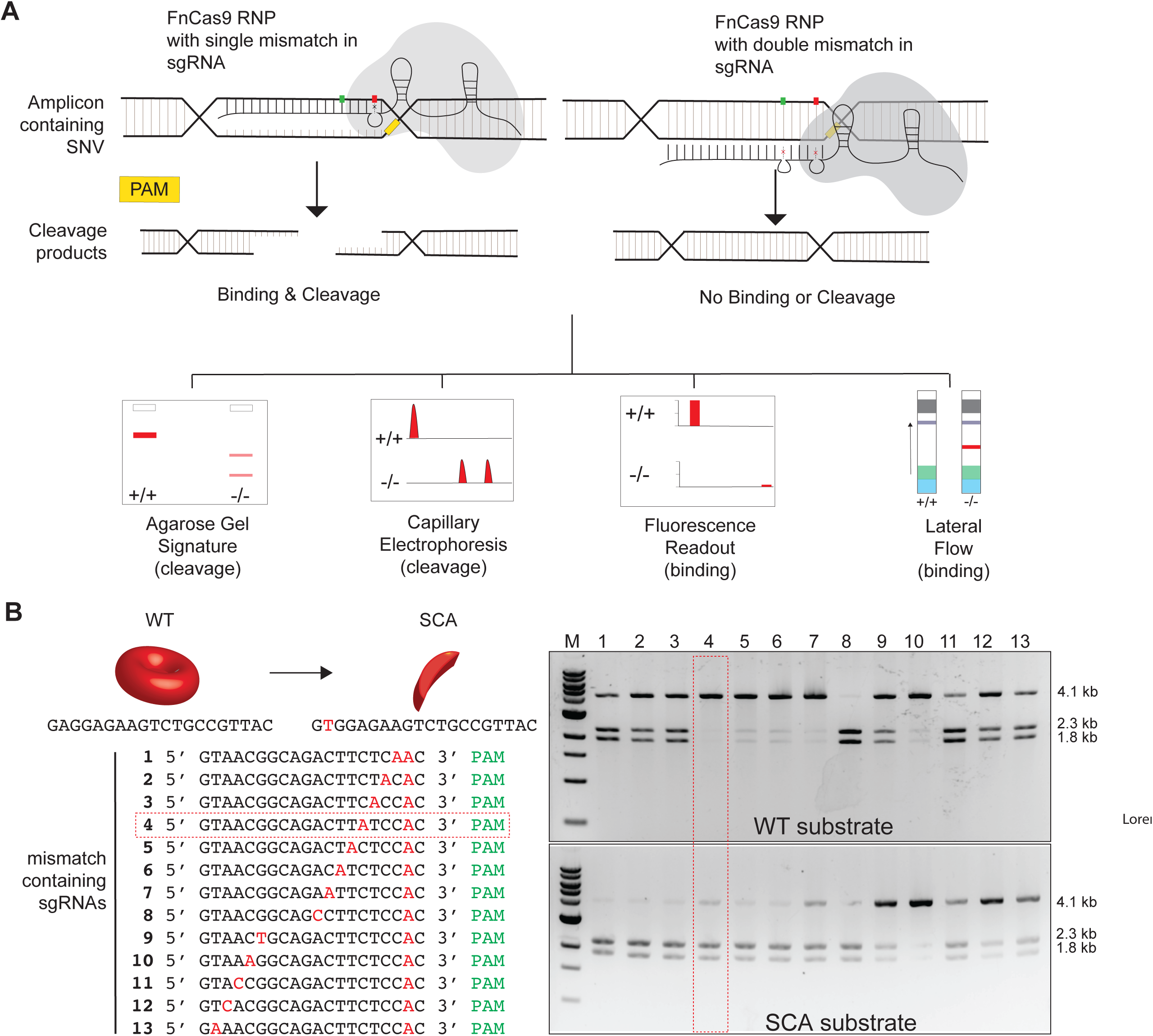
Schematic of FELUDA detection. **A**, Upper panel shows strategy for discrimination of substrates differing by single mismatch using FnCas9. Presence of 2 mismatches (marked in red and green) at defined positions on the sgRNA prevents the enzyme from binding to the target leading to different binding and cleavage outcomes. Readouts can be through gel electrophoresis, capillary electrophoresis, fluorescence measurement or lateral flow assay. **B**, Left panel shows positions of mismatches sgRNAs containing two mismatches at different positions along their lengths. Representative in-vitro cleavage outcomes on wild type (WT) or sickle cell anemia (SCA) substrates (4.1 kb) are shown on the right. Cleavage with FnCas9 produces 2 products (2.3 kb and 1.8 kb). Red dotted box denotes the sgRNA showing negligible cleavage for WT substrate and maximum cleavage for SCA substrate.

To identify a SNV with high accuracy, we first sought to investigate if FnCas9 can be directed to cleave the wild type (WT) variant at a SNV by placing an additional mismatch in the sgRNA sequence specific to the SNV. To test this, we selected sickle cell anemia (SCA), a global autosomal recessive genetic disorder caused by one point mutation (GAG>GTG)^23-24^. We identified an sgRNA that can recognize the main disease causing point mutation (GAG > GTG) on account of a PAM site in the vicinity (Figure 1B). We then fixed the position of this mutation with respect to PAM and changed every other base in the sgRNA sequence to identify which combination led to complete loss of cleavage of a wild type substrate in an *in vitro* cleavage (IVC) assay with FnCas9 (Figure 1B, Supplementary Figure 1A). We found that two mismatches at the 2^nd^ and 6^th^ positions away from the PAM completely abrogated the cleavage of the target (Figure 1B). Similarly other combinations (2 and 7, 2 and 8, 2 and 9, and 2 and 12, numbers referring to positions away from PAM) also abolished the cleavage, although to slightly lower levels (Figure 1B). Notably, on the SCA substrate, the 2 and 6 mismatch combination produced near complete cleavage suggesting that this combination may be favorable for discriminating between WT and SCA substrates (Figure 1B).

Importantly, the same design principle can guide the allelic discrimination at every SNV that appears in these positions upstream of the trinucleotide NGG PAM in DNA. We performed FELUDA with 4 synthetic sequences corresponding to SNVs reported for the Mendelian disorders like Glanzman’s Thrombasthenia, Hemophilia A (Factor VIII deficiency), Glycogen Storage Disease Type I and X linked myotubular myopathy and observed an identical pattern of successful discrimination at the SNVs (Supplementary Figure 1B). Taken together, these experiments suggest that FELUDA design can be universally used for detection of SNVs and and would not require extensive optimization or validation steps for new SNVs. To aid users for quick design and implementation of FELUDA for a target SNV, we have developed a webtool JATAYU (Junction for Analysis and Target Design for Your FELUDA assay) that incorporates the above features and generates primer sequences for amplicon and sgRNA synthesis (https://jatayu.igib.res.in, Supplementary Figure 2).

We next tested FELUDA in DNA from 6 SCA patients and a healthy control and found that in every case, the SC mutation containing substrates were cleaved to give a distinguishable signature while the WT substrate remained intact (Supplementary Figure 3A). To establish that enzyme specificity for position-specific mismatches with respect to PAM site is the fundamental reason for this discrimination, we designed the mismatch combinations such that cleavage will occur only for the WT substrate and observed identical results thus confirming our hypothesis (Supplementary Figure 3B). Notably, a simple agarose gel electrophoresis can be employed for this discrimination suggesting that FELUDA can be used in routine molecular biology labs to establish the presence of an SNV in a DNA sequence.

In recent times fluorescence based nucleic acid detection has been widely used for several CRISPRDx platforms, particularly where collateral cleavage of reporters has been employed. Although FELUDA results can be precisely determined by agarose or capillary electrophoresis, we envisioned fluorescence or chemiluminescence as alternate end-point readouts to expand the scope of devices that can suit FELUDA based detection. To implement this, we first investigated if FELUDA can be adapted to a non-cleavage, affinity-based method of detection which works with single nucleotide mismatch sensitivity.

To develop such a readout, we tested FELUDA with a catalytically dead FnCas9 (dFnCas9) tagged with a fluorophore (GFP) and investigated if its mismatch discrimination is regulated at the level of DNA binding (Supplementary Figure 1A). We performed microscale thermophoresis (MST) assays to measure the binding affinity of inactive FnCas9-GFP RNP complex with WT or SCD substrates. We observed that the SCD substrate showed moderately strong binding (K_d_ = 187.2 ± 3.4 nM) whereas the WT substrate exhibited very weak binding (K_d_ = 1037.4 nM ± 93.3 nM) consistent with the absence of cleavage on the IVC readout (Figure 2A). We then developed a pipeline to adapt FELUDA for an affinity-based fluorescent read-out system, where the amplification step generates biotinylated products which can then be immobilized on magnetic streptavidin beads. Upon incubation with fluorescent components in FELUDA, the absence or presence of a mismatch guides the binding of FnCas9 molecules to the substrate leading to loss of fluorescence signal in the supernatant (Figure 2B). We tested FELUDA using WT or SCA substrates and observed distinct signatures that distinguished between the two alleles suggesting that FELUDA can be adapted for a fluorescence-based readout (Figure 2B).

**Fig. 2:**
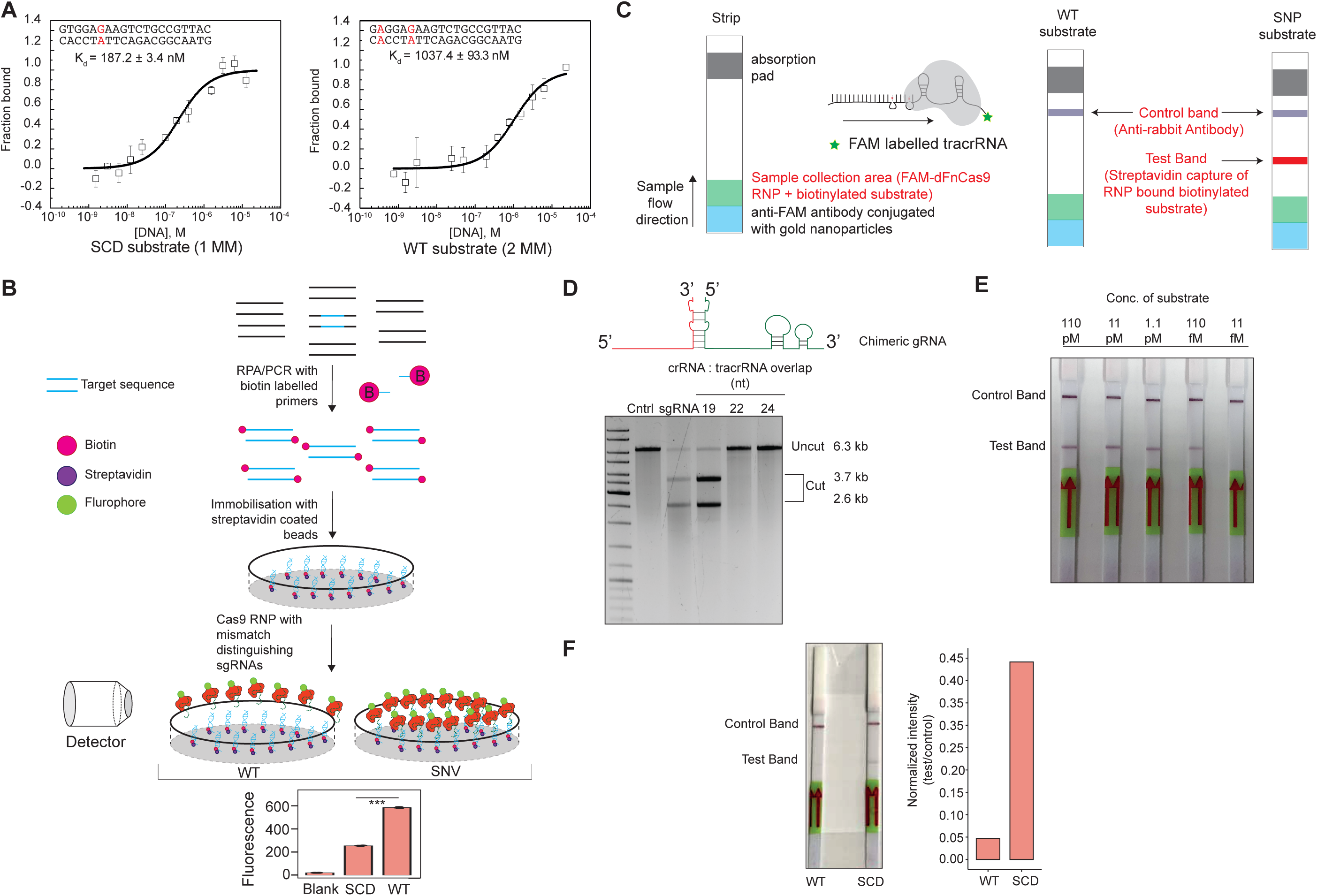
FELUDA can be adapted to multiple readout modes. **A**, Binding experiments using Microscale Thermophoresis showing interaction of FnCas9 with substrates with 1 mismatch (MM) on the left and 2 mismatches on the right. Values are expressed as fraction bound protein (y-axis) with respect to varying concentrations of purified DNA substrate (Molar units, M, x-axis). Error bars represent SEM (2 independent experiments). **B**, Schematic for fluorescence based detection of sickle cell anemia mutation using FELUDA. Error bars represent SD (3 independent experiments). **C**, Outline of lateral flow assay using FELUDA showing positions of control and test bands. **D**, Adaptation of chimeric gRNA for FELUDA. Cleavage outcomes with different lengths of overlap between crRNA and tracrRNA are shown. **E**, Detection of varying concentrations of substrate DNA (HBB) using FELUDA (pM = picomolar, fM = femtomolar). **F**, Detection of SCA mutation using FELUDA on a paper strip. Quantification of band intensities shown on the right.

Next, we investigated if FELUDA detection can be extended to a quick point of care diagnosis of SNVs using lateral flow strips for instrument-free visual detection. Although such read-outs have been demonstrated with CRISPR effectors that have collateral activity, they rely on a secondary signal amplification step where fluorescent oligos are added to the reaction setup. We designed an assay using FAM labelled RNP complex and biotin labelled amplicons on commercially available paper strips. In such a reaction, RNP-bound substrate molecules in a solution can lead to aggregation of the complex on a distinct test line of the strip while anti-FAM antibody linked gold nanoparticles accumulate on a control line on the strip (Figure 2C). To enable FAM labelling of sgRNA, we first validated the successful activity of chimeric FnCas9 sgRNAs by altering the length of overlap between crRNA:tracrRNA and observed that 19nt overlap can efficiently cleave the target (Figure 2D). Since a single FAM labelled tracrRNA is compatible with multiple crRNAs, this design also reduces the time and cost of a FELUDA assay. Next, we performed this assay with the HBB target and were able to detect up to femtomolar levels of target DNA in a solution suggesting that even without signal amplification, FELUDA reaches sensitivity similar to that reported for CRISPRDx platforms employing collateral activity (Figure 2E). Finally, we tested FELUDA using WT and SCA samples and obtained clear distinguishing bands for either condition validating the feasibility of visual detection for FELUDA diagnostics with complete accuracy (Figure 2F).

Genotyping carrier individuals with heterozygosity though non-sequencing PCR based methods is often complicated and requires extensive optimization of primer concentration and assay conditions. Although sickle cell trait (SCT) individuals are generally non-symptomatic, carrier screening is vital to prevent the spread of SCA in successive generations and is widely employed in SCA control programs in various parts of the world^19^. We speculated that the high specificity of FELUDA can be extended to identifying carriers since FnCas9 would cleave 50% of the DNA copies carrying SCA mutation and thus show an intermediate cleavage pattern (Figure 3A.) We also investigated the use of saliva instead of blood as a non-invasive source of genomic DNA that would allow genotyping children and aged subjects where drawing blood may not be feasible. We obtained a clear, distinguishable signature of DNA cleavage in an SCT subject that was intermediate between normal or SCA individuals suggesting that FELUDA can be successfully used for determining zygosity at a specific SNV (Figure 3A). Although this reinforced the inherent sensitivity of FELUDA for genotyping targets with 1bp mismatches, obtaining the three distinct readouts would necessitate significantly robust reaction components and high reproducibility across multiple subjects. We performed a blinded experiment using DNA obtained from 49 subjects with all three genotypes from a Tertiary care center. Remarkably, the FELUDA results perfectly matched with the sequencing data from the same samples performed in a different laboratory (CSIR Center for Cellular and Molecular Biology, Chandak Lab) and thus identified all three genotypes with 100% accuracy (Figure 3B).

**Fig. 3:**
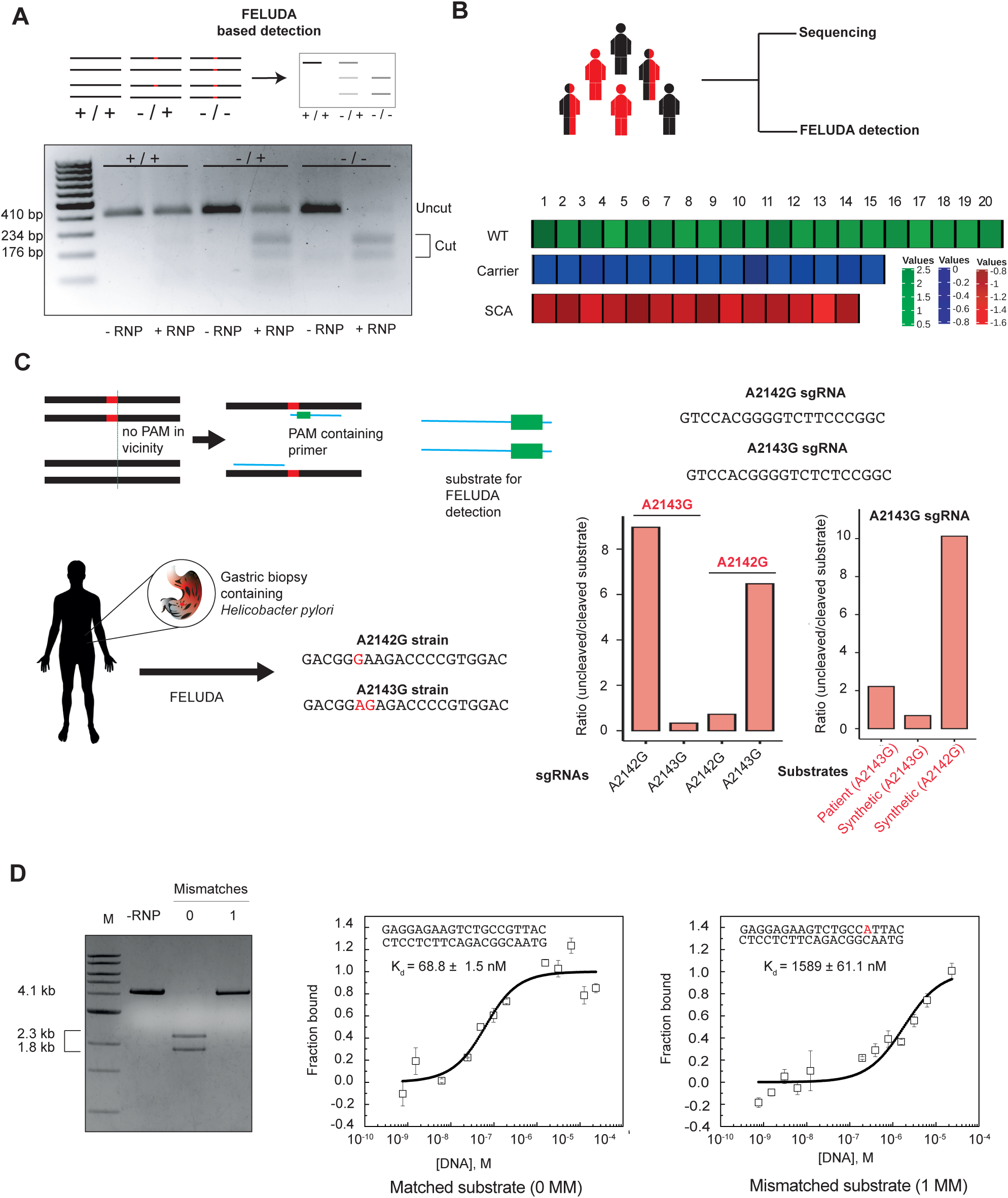
Detection of zygosity and PAM independent FELUDA design. **A**, Schematic of FELUDA for identifying carriers of SCA mutation. Representative gel image shows distinct FELUDA signatures for Wild type (WT), Carrier or SCA substrates. **B**, FELUDA accurately identifies zygosity in a mixed cohort of individuals. Each subject is color coded according to z-scores of their gel cleavage outcomes (uncleaved/cleaved). **C**, Top, strategy for detection of SNVs with an in-built PAM (PAM-mer) in primer sequence. Bottom, closely related strains of *H. Pylori* showing the mutations they are associated with in red. Right, two different sgRNAs, A2142G and A2143G can produce distinct cleavage outcomes with their respective substrates (shown in red, bottom left). Bottom right shows FELUDA based detection of A2143G genotype of *H Pylori* from gut biopsy of a patient. Cleavage outcomes with synthetic substrates for A2143G and and A2142G are also shown. **D**, Representative gel image showing no cleavage of HBB substrate with sgRNA containing 1 mismatch (16^th^ position, PAM distal). Binding experiments using Microscale Thermophoresis showing high binding affinity with no mismatches to substrate and negligible binding affinity when 1 mismatch at 16^th^ position (PAM distal) is present (MM, mismatches). Values are expressed as fraction bound protein (y-axis) with respect to varying concentrations of purified DNA substrate (Molar units, M, x-axis). Error bars represent SEM (2 independent experiments).

Although several pathogenic SNVs are located close to a NGG PAM and can be accurately targeted by FELUDA, these form only a small subset of the total number of disease-causing SNVs catalogued in ClinVar database (Supplementary Figure 4)^25^. To detect the non-PAM proximal SNVs we designed an in-built PAM site in the amplification step of FELUDA. We tested this approach using 2 SNVs (A2142G and A2143G) present in *Helicobacter pylori* 23s *rRNA* gene and which do not have a NGG PAM in the vicinity. These SNVS confer variable clarithromycin resistance in patients with gastric ulcers and clinically pose a serious concern for physicians^26^. We validated that PAM-mer based amplification can be successfully used for FnCas9 based IVC by targeting synthetic DNA sequence containing the *H. pylori* 23s *rRNA* gene (Figure 3C). We then isolated bacteria DNA from patient gut biopsy samples and successfully distinguished the antibiotic resistance genotype from another closely matched synthetic wild type sequence (Figure 3C). Importantly, this procedure takes a few hours from obtaining sample to diagnosing the variant, a significant improvement over existing regimens which rely on antimicrobial susceptibility tests that can take several days to complete^27^.

Since our design incorporates the need for fixing the SNV at either 2^nd^ or 6^th^ position proximal to PAM, this limits the possibility of discriminating substrates using a single FELUDA reaction. To overcome this, we explored single mismatches in the sgRNA sequence that might lead to abrogation of cleavage or binding from the substrate particularly at positions 16-19 at PAM distal end, since these showed minimum cleavage from our previous study. Remarkably, we found that FnCas9 shows negligible cleavage at each of these positions (Supplementary Figure 5A). In particular, mismatch at PAM distal 16^th^ base shows complete absence of cleavage and negligible binding affinity to mismatched substrate (Figure 3D). To confirm this strategy, we targeted the SNV rs713598 which is associated with either G or C or G/C (heterozygous) variants in different individuals^28^. Using our single base pair mismatch approach we could successfully distinguish individuals with each of these genotypes (Supplementary Figure 5B).

FELUDA detection can be customized with other features of current CRISPRDx methods such as detecting SNVs in RNA or compatibility with isothermal amplification of substrate^29^. At the level of RNA, by appending a Reverse Transcription (RT) step prior to amplification, we could successfully distinguish two Dengue virus serotypes (West Pac/74 and P23085, designated 1 and 2) that lead to variable mortality rates in patients (Supplementary Figure 6)^30,31^. To circumvent the need of complex instrumentation for preamplification, we optimized Recombinase Polymerase Amplification (RPA) for generating amplicons that can contribute to a field deployable assay prototype (Supplementary Figure 7A-B)^32^. Importantly, FELUDA detection can occur in a wide temperature range (10 °C – 50 °C) and up to 3 days post thawing (at room temperature) suggesting that field studies using FELUDA can be conducted in diverse climatic conditions and reaction components can be successfully used following cold chain transportation (Supplementary Figure 8A-B).

During the preparation of this manuscript, a sudden outbreak of Corona virus disease 19 (COVID-19) due to SARS-CoV-2 virus infection broke out and has led to an ongoing public health emergency in most countries of the world. In addition to general social distancing, identification of infected individuals and screening their contacts for possible quarantine measures is one of the major steps in reducing community transmission of the virus^33^. It is postulated that rapid and accurate testing protocols can significantly aid in this process^34^. We explored FELUDA as a simple alternative to quantitative PCR based detection of SARS-CoV-2 virus since it does not need complex instrumentation (Figure 4A). We designed sgRNAs targeting PAM containing conserved regions (*NSP8* and Nucleocapsid phosphoprotein) across all strains in the SARS-CoV-2 viral genome (229 sequences deposited till 30^th^ March 2020) and sgRNAs targeting conserved regions across the Influenza virus H1N1 genome (45 complete genome sets deposited till 15^th^ March 2020) since both infections have overlapping symptoms. Additionally, we selected human *ACTB* as a positive control for human DNA traces that typically accompany the viral DNA extracted from nasal or buccal swabs. Using FELUDA, we obtained clear signatures of SARS-CoV-2 sequence in synthetic DNA using a specific RNP that is distinguishable from non-specific RNP (such as *H1N1* or *HBB*) (Figure 4B). Importantly FELUDA was also able to distinguish between two SARS-CoV-2 and SARS-CoV-1 sequences that differ by a single nucleotide (Supplementary Figure 9A)^35^. Remarkably, FELUDA based lateral flow assay could distinguish SARS-CoV-2 synthetic DNA on a paper strip only when SARS-CoV-2 specific RNP was used to interrogate the substrate (Figure 4C). Finally, we confirmed the applicability of such lateral flow device as a rapid, cost effective and machine-independent alternative to current testing protocols by successfully detecting the presence of viral signature from low amounts of total RNA obtained from COVID-19 patients within one hour (Figure 4D, Supplementary Figure 9B). We have compiled a standardized protocol for performing this test on clinical samples (Supplementary Note 1).

**Fig. 4:**
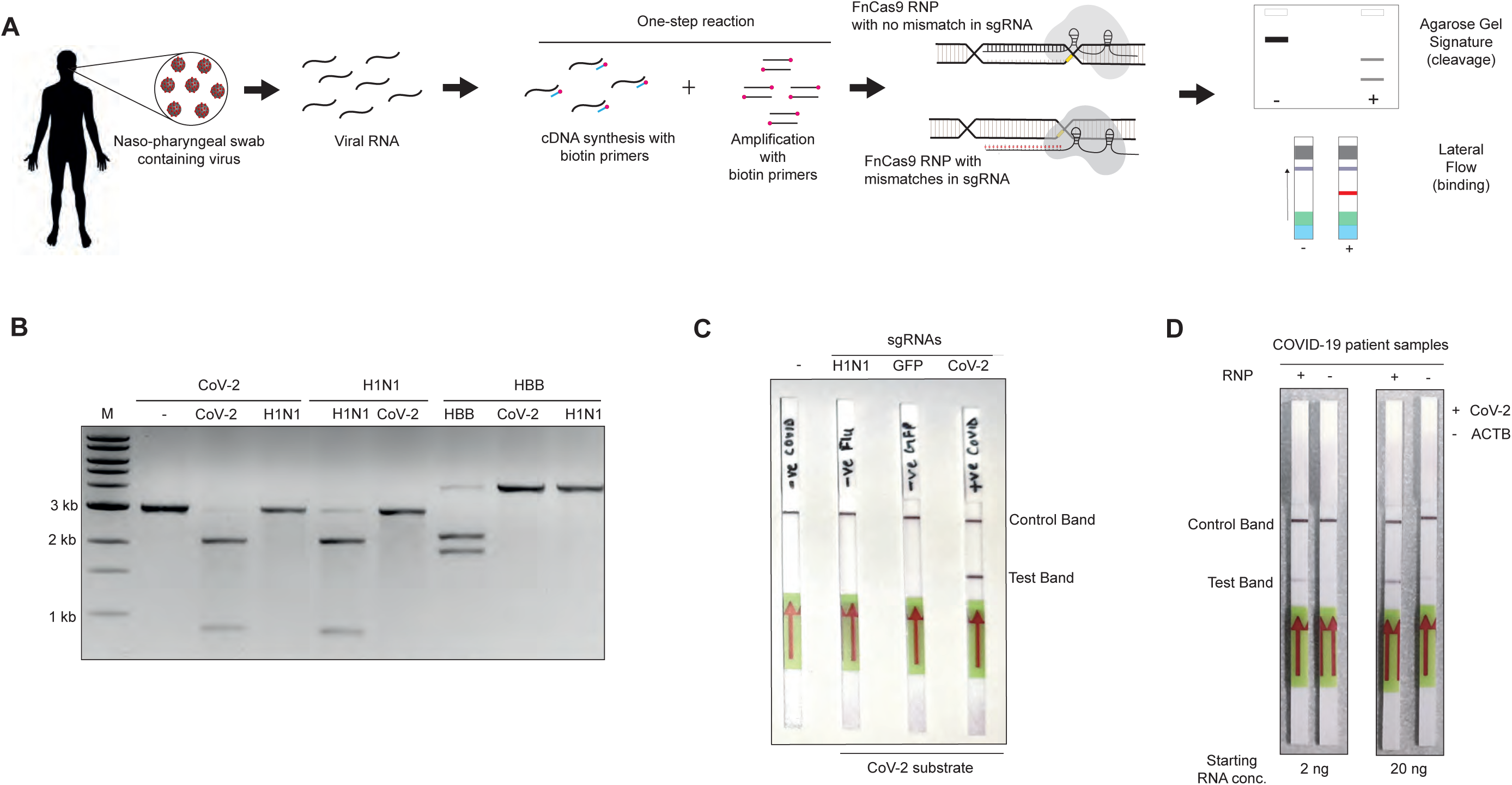
Deployment of FELUDA for rapid and accurate diagnosis during COVID-19 pandemic. **A**, Pipeline of FELUDA based detection for SARS-CoV-2 infection in samples obtained from patients. Individual steps are depicted. **B**, Successful identification of SARS-CoV-2 (COVID), H1N1 yeand HBB substrates with respective sgRNAs. **C**, CoV-2 specific FELUDA can detect CoV2 target on paper strip but non-specific RNPs do not produce detectable signal. **D**, Specific detection of CoV-2 in COVID-19 patient samples (2 ng and 20 ng starting RNA) using FELUDA. ACTB serves as control.

In this report, we present the versatile applications of FELUDA for rapid and accurate detection of nucleic acids at a low cost (Supplementary Figure 9C-D). Leveraging the highly specific binding and subsequent cleavage properties of FnCas9 as distinct enzymatic steps, FELUDA couples sensitivity with broad spectrum of read-out possibilities that can be carried out in the lab or in the field. Detection of low copy numbers of nucleic acids usually involves signal amplification through collateral cleavage of reporter molecules on other CRISPRDx platforms but FELUDA is highly accurate in diagnosing SNVs without them. Its strength lies in unbiased identification of nucleobases and their variants. The affinity based FELUDA assays can be used for designing panels of mutation-scanning sgRNAs on a microchip that can give rise to rapid readouts from patient samples for multiple targets simultaneously (Supplementary Figure 10A). The combination of single mismatch sensitivity and the possibility of tagging different fluorophores to the RNP can allow identifying each of the 4 nucleobases in a target at a given position. This can potentially expand the applications of CRISPR in the domain of affinity based target sequencing (Supplementary Figure 10B). Its ease of design and implementation, as exemplified by its urgent deployment during the COVID-19 health crisis offers immense possibilities for rapid and wide-spread testing that has so far proven to be successful in spreading the progression of the disease in multiple countries.

## Methods

### Plasmid Construction

The sequences for the Hbb (WT and SCA), EMX1 and VEGFA site 3 were PCR amplified and cloned into the TOPO-TA vector (Thermo Fisher Scientific). Around region of around 500bp including SNV/mismatches for four Mendelian disorders Glanzman, Thrombasthenia, Hemophilia A (Factor VIII deficiency), Glycogen Storage Disease Type I and X-linked myotubular myopathy were procured as synthetic DNA oligos and cloned into the pUC57 by EcoRV (GENESCRIPT). Similarly, a 500bp region flanking two SNVs (A2142G and A2143G) in *Helicobacter pylori* 23s *rRNA* gene were ordered as synthetic DNA cloned in pUC57 by EcoRV. The cloned sequences were confirmed by Sanger sequencing.

### Protein purification

All the plasmid constructs used were adapted from our previous study^16^. Briefly, *Escherichia coli* Rosetta2 (DE3) (Novagen) were used to express the protein (FnCas9-WT, dFnCas9 and dFnCas9-GFP) which was induced by 0.5 mM isopropyl β-D-thiogalactopyranoside (IPTG) and cultured at 18°C for overnight. The cell lysate was first purified using Ni-NTA beads (Roche) and then subjected to size-exclusion chromatography on HiLoad Superdex 200 16/600 column (GE Healthcare) in 20 mM HEPES pH 7.5, 150 mM KCl, 10% glycerol, 1mM DTT and 10mM MgCl_2._ The concentration of purified proteins were measured by Pierce BCA protein assay kit (Thermo Fisher Scientific). The purified proteins were stored at −80°C until further use.

### *In vitro* transcription (IVT)

*In vitro* transcription for sgRNAs/crRNAs were done using MegaScript T7 Transcription kit (Thermo Fisher Scientific) using a T7 promoter containing template as substrates. IVT reactions were incubated overnight at 37°C followed by DNase treatment as per kit instructions and then purified by NucAway spin column (Thermo Fisher Scientific) purification. IVT sgRNAs/crRNAs were stored at −20°C until further use.

### Human Genomic DNA extraction

Human Genomic DNA was extracted from the blood using the Wizard Genomic DNA Purification kit (Promega) as per the instructions.

For genomic DNA extraction from saliva, 1ml of saliva was centrifuged at 13000 rpm followed by three washes with 1ml of 1X PBS. After washing, the pellet was lysed with 50µl of 0.2% Triton X100 at 95°C for 5 minutes. Then again centrifuged at 13000 rpm and supernatant was transferred into a fresh vial. A total volume of 1µl of the supernatant was used in PCR reaction or otherwise stored at −20 °C.

### Bacterial genomic DNA extraction

Genomic DNA was extracted from the biopsy samples (15-20 mg) of patients infected with *Helicobacter Pylori* using DNeasy 96 PowerSoil Pro QIAcube HT Kit using Vortex Genei for the sample lysis. The sample was eluted in 100µl of nuclease free water using a vacuum pump.

### Polymerase Chain Reaction (PCR)

DNA sequences containing mutations associated with normal of sickle anemia mutation within Hbb locus were were amplified. For PCR, 50 ng of the genomic DNA extracted from the wild type and the sickle cell patient’s saliva or blood in 50 µl PCR reaction containing 500 nM of Forward and Reverse Primer each (normal or biotinylated), 200 nM of dNTPs, 1X reaction buffer with the Taq DNA polymerase were used.

### Recombinase Polymerase Amplification (RPA)

RPA reaction was set up as recommended in the TwistAmp^®^ Basic kit. 3-5 ng of genomic DNA was used with normal or biotinylated primers and the reaction was performed at 39°C for 20 minutes. Amplicons were then purified with Qiagen PCR clean up kit and visualized on agarose gel.

### Gel Based Nucleic Acid Detection

#### *In vitro* Cleavage assay (IVC)

DNA samples were directly PCR amplified for the target region while RNA samples were converted to cDNA using Reverse Transcriptase kit (Qiagen) and PCR amplified. In vitro cleavage assay was performed as optimized in our previous study^16^. For in vitro cleavage, 100ng of cleaned amplicon or linearized plasmid was incubated in reaction buffer (20 mM HEPES, pH7.5, 150mM KCl, 1mM DTT, 10% glycerol, 10mM MgCl_2_) along with reconstituted RNP complex (500nM) at 37°C for 30 minutes and cleaved products were visualized on agarose gel.

#### Fluorescence Detection

DNA regions from *HBB* locus (wild type and Sickle cell sample) were amplified using biotinylated primers to biotinylate the amplicons. dFnCas9-GFP: sgRNA (180 nM:540 nM) RNP complex was reconstituted at 25°C for 10 mins in reaction buffer (20 mM HEPES, pH7.5, 150mM KCl, 1mM DTT, 10% glycerol, 10mM MgCl2). Meanwhile, 6µL of the Dynabeads MyOne Streptavidin C1 (Thermo Fisher Scientific) were washed three times with 30µl of buffer containing 50 mM Tris-Cl (pH7.5), 5 mM EDTA, 1M NaCl. Then washed beads were incubated with 1µM of biotinylated Hbb amplicon (wild type or Sickle cell sample) for 30 minutes in the reaction buffer. The reconstituted RNP complex was incubated with the streptavidin bound PCR amplicon for 30 minutes. Further, emission spectra of unbound dFnCas9-GFP was measured using Monolith NT. 115 (NanoTemper Technologies GmbH, Munich, Germany) where sample was loaded into standard treated capillaries provided by Nanotemper. Measurements were performed using 60% excitation power in blue filter (465-490nm excitation wavelength; 500-550nm emission wavelength) with medium MST power.

#### Lateral Flow Detection

Chimeric gRNA (crRNA:TracrRNA) was prepared by mixing *in-vitro* transcribed crRNA and synthetic 3’-FAM labelled TracrRNA in a equimolar ratio using annealing buffer (100mM NaCl, 50mM Tris-HCl, pH 8.0 and 1mM MgCl_2_), mix was heated at 95CC for 2-5 minutes and then allowed to cool at room temperature for 15-20 minutes. Chimeric gRNA-dead FnCas9 RNP complex (500nM) was prepared by mixing them in a buffer (20mM HEPES, pH7.5, 150mM KCl, 1mM DTT, 10% glycerol, 10mM MgCl_2_) and incubated for 10 min at RT. RPA or PCR amplified biotinylated amplicons were then incubated with the RNP complexes for 15 minutes at 37°C. Dipstick buffer was added along with the Milenia Hybridetect paper strip (TwistDx) was added as per instructions, producing dark colored bands over the strip within 2-10 min.

#### FnCas9 activity in a temperature range

FnCas9-sgRNA complex (500nM) was prepared by mixing them in a buffer (20mM HEPES, pH7.5, 150mM KCl, 1mM DTT, 10% glycerol, 10mM MgCl_2_) and incubated for 10 min at RT. The reconstituted RNP complexes along with PCR amplified DNA amplicons were then used for IVC assays at different temperatures ranging from **10°C to 50°C** for 30 minutes. The reaction was inhibited using 1µl of Proteinase K (Ambion). After removing residual RNA by RNase A (Purelink), the cleaved products were visualized on agarose gel.

#### Stability of the FnCas9 protein

FnCas9-sgRNA complex (500nM) was prepared by mixing them in a buffer (20mM HEPES, pH7.5, 150mM KCl, 1mM DTT, 10% glycerol, 10mM MgCl_2_) and incubated for 10 min at RT. Linearized plasmid used here as a substrate was incubated with reconstituted RNP complexes at different time points starting from 0 h to 100 h with or without 10% sucrose in the reaction buffer respectively. Further, cleaved products were visualized on agarose gel.

#### MST Method

For the binding experiment, dFnCas9-GFP protein was used along with PAGE purified respective IVT sgRNAs. Notably, IVT sgRNAs were purified by 12% Urea-PAGE. The binding affinities of the dFnCas9 protein and sgRNA (ribonucleoprotein) complexes with targets were calculated using Monolith NT. 115 (NanoTemper Technologies GmbH, Munich, Germany). RNP complex (Protein:sgRNA molar ratio,1:1) was formed at 25 °C for 10 mins in reaction buffer (20 mM HEPES, pH7.5, 150mM KCl, 1mM DTT, 10mM MgCl_2_). For the target DNA constitution, duplex was formed using HPLC purified 30bp ssDNA oligo (Sigma) by annealing for 5 mins at 95 °C and then by slowly cooling down at 25 °C. 30 bp dsDNA of different genomic targets with varying concentrations (ranging from 0.7nM to 25μM) were incubated with RNP complex at 37^°^ C for 60 min in reaction buffer. NanoTemper standard treated capillaries were used for loading the sample. Measurements were performed at 25°C using 40% LED power in blue filter (465-490nm excitation wavelength; 500-550nm emission wavelength) and 40% MST power. All experiments were repeated at least three times for each measurement. All Data analyses were done using NanoTemper analysis software. The graphs were plotted using OriginPro 8.5 software.

#### Sanger Sequencing

The sequencing reaction was carried out using Big dye Terminator v3.1 cycle sequencing kit (ABI, 4337454) in 10μl volume (containing 0.5μl purified DNA, 0.8μl sequencing reaction mix, 2μl 5X dilution buffer and 0.6μl forward/reverse primer) with the following cycling conditions - 3 mins at 95°C, 40 cycles of (10 sec at 95°C, 10 sec at 55°C, 4 mins at 60°C) and 10 mins at 4°C. Subsequently, the PCR product was purified by mixing with 12μl of 125mM EDTA (pH 8.0) and incubating at RT for 5 mins. 50μl of absolute ethanol and 2μl of 3M NaOAc (pH 4.8) were then added, incubated at RT for 10 mins and centrifuged at 3800rpm for 30 mins, followed by invert spin at <300rpm to discard the supernatant. The pellet was washed twice with 100μl of 70% ethanol at 4000rpm for 15 mins and supernatant was discarded by invert spin. The pellet was air dried, dissolved in 12μl of Hi-Di formamide (Thermo fisher, 4311320), denatured at 95°C for 5 mins followed by snapchill, and linked to ABI 3130xl sequencer. Base calling was carried out using sequencing analysis software (v5.3.1) (ABI, US) and sequence was analyzed using Chromas v2.6.5 (Technelysium, Australia).

#### Disease selection from ClinVar Database

ClinVar dataset (version: 20180930) was used to extract disease variation spectrum that can be targeted by FELUDA^25,36^. Only Single nucleotide variations (SNVs) were analyzed. Further, SNVs which are situated 2 bp upstream of the PAM sequence were extracted. Next, SNVs with Pathogenic effects and valid OMIM ID were used for further analysis. Finally, variations with higher frequency in Indian Population were selected for the validation. Custom python script was used for the whole process.

#### sgRNA design for CoV-2, CoV-1 and H1N1

To differentiate between CoV-2 (MN908947.3) and CoV-1 (NC_004718.3) sequences^37-38^ the FELUDA pipeline with 16^th^ base (PAM distal) mismatch was followed. Overall, 5 sgRNAs were extracted which show different single nucleotide variation between CoV-2 and CoV-1. To find out sgRNAs specific to strain/region, 229 sequences (Gene sequence and full genome) of CoV-2 were extracted and custom python script and SeqMap were used to design strain specific sgRNAs. For H1N1, Influenza Virus Databases were used^39^. Sequences of virus with human host and Indian region were extracted. Overall, 45 genomes set with 360 sequences were downloaded. Next, the common sgRNA among all the sequences were fetched using custom python script and SeqMap^40^.

#### JATAYU

JATAYU (jatayu.igib.res.in) is a web tool which enables users to design sgRNAs and primer for the detection of variations. Users need to provide a valid genomic sequence with position and type of variation. JATAYU front-end has been created using Bootstrap 4 and jQuery. In the back-end, python-based Flask framework has been used with genome analysis tools BWA (Burrows-Wheeler aligner) and bedtools^41-42^.

All oligos used in the study are listed in Table 1.

## Acknowledgments

We thank all members of Chakraborty and Maiti labs for helpful discussions and valuable insights. We are grateful to Mitali Mukerji, Rajesh Pandey and Mohd. Faruq (CSIR IGIB) and Sankar Bhattacharyya (Translational Health Science and Technology Institute, Faridabad, India) for providing DNA and RNA samples used in the study. This study was funded by CSIR Sickle Cell Anemia Mission (HCP0008) and a Lady Tata Young Investigator award to D.C.

## Contributions

D.C., S.M., M.A. and R.P. conceived, designed and interpreted the experiments. A.H.A provided bioinformatics support. D.S. performed MST experiments. N.S. performed mismatch-based cleavage assays. M.K. performed assays on COVID-19 detection. M.Ai. and S.S. performed studies on discriminating single nucleotide variants using FELUDA. G.M. designed studies on detecting *H Pylori* variants. H.L. and P.K.P. helped in the random screening of school children and identifying sickle cell anemia patients at Chhattisgarh. G.R.C performed sequencing of SCA patient samples. D.C. wrote the manuscript with inputs from S.M. and G.R.C.

## Ethics declarations

The present study was approved by the Ethics Committee, Institute of Genomics and Integrative Biology, New Delhi. D.C., S.M., M.A., R.P., A.S.A, D.S., N.S. and S.S. are authors in provisional patent applications that have been filed in relation to this work.

## Supplementary Figure Legends

**Supplementary Figure 1: FELUDA can successfully discriminate pathogenic SNVs. A**, Representative Coomassie gel showing proteins used in the study. **B**, FELUDA design can universally discriminate SNVs across diverse targets. Representative gel image showing synthetic targets containing either WT or pathogenic SNVs. FELUDA can distinguish between the two variants.

**Supplementary Figure 2: Schematic of JATAYU**. Main steps in the pipeline are shown. Step 1: Users can enter the sequence with length from 20 up to 30 nucleotides.

Step 2: Users need to provide mutation information such as Position and type of mutation. Step 3: Confirmation of the mutation information. Step 4: Design of sgRNA and primers for the given input sequence and mutation.

**Supplementary Figure 3: FELUDA can successfully distinguish SCA mutation from patient derived DNA. A**, Representative agarose gel electrophoresis showing cleavage of 6 SCA patient derived DNA sequences (2-7) using an sgRNA with 2 mismatches to wild type sequence. Healthy individual’s DNA used as control (1). **B**, Cleavage outcomes can be swapped by placing 2 mismatches to SCA sequence and is shown on a representative agarose gel.

**Supplementary Figure 4: NGG PAM vs Non-NGG PAM targets for FELUDA:** Only a small fraction of pathogenic variants catalogued in ClinVar dataset can be targeted by conventional FELUDA. Small subset (in red) in every pie chart belongs to targetable mutation with NGG PAM and bigger set in different colors represent non NGG PAM mutation targets.

**Supplementary Figure 5: FELUDA can successfully distinguish substrates on the basis of 1 mismatch. A**, Representative agarose gel showing GFP substrate cleaved with sgRNAs harboring mismatches at PAM distal 16-19^th^ position. WT sgRNA (0) shows successful cleavage. **B**, Top, Circos plot showing the location of the allelic variant rs713598 on the human genome. Bottom, Sanger sequencing confirming the allelic variants obtained from 4 subjects and FELUDA design based on placing mismatch at 16^th^ position enables detection of individual variants. Negligible cleavage is seen with G variant, maximum cleavage is seen with C variant and intermediate cleavage seen with G/C variant.

**Supplementary Figure 6: FELUDA can distinguish SNVs from RNA samples**. Pipeline of detection from RNA samples is shown, including a Reverse Transcription step. On the right, representative gel image of 2 Dengi Virus variants (1 and 2) differing by a single mismatch is shown. FELUDA design produces variant-specific agarose gel signatures.

**Supplementary Figure 7: FELUDA is compatible with isothermal amplification of DNA substrates. A**, Representative gel image of EMX1 substrate amplified using Recombinase Polymerase Amplification (39 °C, 20 min) is shown. **B**, In vitro cleavage outcome of RPA product on gel.

**Supplementary Figure 8: FELUDA components are robust across different conditions. A**, Representative gel images shows successful cleavage of substrates at different temperatures (10 °C – 50 °C). **B**, Representative gel images show FnCas9 activity after incubation at room temperature upto 72 h post thawing. Red dotted box shows loss of activity at 100 h.

**Supplementary Figure 9: FELUDA for rapid detection of CoV-2 presence in samples. A**, FELUDA can successfully discriminate between CoV-1 and CoV-2 substrates that differ by 1 mismatch. Representative gel image showing distinct outcomes of FELUDA on individual substrates. **B**, FELUDA can detect a specific target (ACTB) from up to 50 pg of starting RNA. **C**, FELUDA reaction for CoV-2 detection can be done within 1h (45 min with one-step RT-PCR or 32 min with RT-RPA reaction). **D**, Cost for a single FELUDA assay for CoV-2 detection is less than 6 USD.

**Supplementary Figure 10: Applications of FELUDA beyond in vitro diagnosis. A**, A microchip containing immobilized FELUDA RNPs, each specific to an SNV can capture only desired substrates producing distinct signal patterns. This will allow multiplexed detection of SNVs from a sample. **B**, By using 4 different fluorescently tagged RNPs and coupling them with sgRNAs containing single mismatches, the identity of a nucleobase at a given position on a DNA sequence can be determined. Only one fluorescent RNP will bind to the target while the other RNPs will dissociate. The process can be iteratively performed to identify each nucleobase in a sequence.

## References

1. Chertow, D. S. Next-generation diagnostics with CRISPR. Science (80-.). 360, 381LP–382 (2018).

2. Li, Y., Li, S., Wang, J. & Liu, G. CRISPR/Cas Systems towards Next-Generation Biosensing. Trends Biotechnol. 37, 730–743 (2019).

3. Pardee, K. et al. Rapid, Low-Cost Detection of Zika Virus Using Programmable Biomolecular Components. Cell 165, 1255–1266 (2016).

4. Teng, F. et al. CDetection: CRISPR-Cas12b-based DNA detection with sub-attomolar sensitivity and single-base specificity. Genome Biol. 20, 1–7 (2019).

5. Quan, J. et al. FLASH: a next-generation CRISPR diagnostic for multiplexed detection of antimicrobial resistance sequences. Nucleic Acids Res. 47, e83 (2019).

6. Kellner, M. J., Koob, J. G., Gootenberg, J. S., Abudayyeh, O. O. & Zhang, F. SHERLOCK: nucleic acid detection with CRISPR nucleases. Nat. Protoc. 14, 2986–3012 (2019).

7. Li, L. et al. HOLMESv2: A CRISPR-Cas12b-Assisted Platform for Nucleic Acid Detection and DNA Methylation Quantitation. ACS Synth. Biol. 8, 2228–2237 (2019).

8. Gootenberg, J. S. et al. Multiplexed and portable nucleic acid detection platform with Cas13, Cas12a, and Csm6. Science (80-.). 360, 439 LP–444 (2018).

9. Sashital, D. G. Pathogen detection in the CRISPR-Cas era. Genome Med. 10, 1–4 (2018).

10. Qiu, X. Y. et al. Highly Effective and Low-Cost MicroRNA Detection with CRISPR-Cas9. ACS Synth. Biol. 7, 807–813 (2018).

11. Abudayyeh, O. O. et al. C2c2 is a single-component programmable RNA-guided RNA-targeting CRISPR effector. Science (80-.). 353, aaf5573 (2016).

12. Li, S. Y. et al. CRISPR-Cas12a has both cis- and trans-cleavage activities on single-stranded DNA. Cell Res. 28, 491–493 (2018).

13. Chen, J. S. et al. CRISPR-Cas12a target binding unleashes indiscriminate single-stranded DNase activity. Science (80-.). 360, 436–439 (2018).

14. Csx, P. et al. Cas13b Is a Type VI-B CRISPR-Associated RNA-Guided RNase Differentially Regulated by Accessory Article Cas13b Is a Type VI-B CRISPR-Associated RNA-Guided RNase Differentially Regulated by Accessory Proteins Csx27 and Csx28. Mol. Cell 65, 618-630.e7 (2017).

15. Yan, W. X. et al. Cas13d Is a Compact RNA-Targeting Type VI CRISPR Effector Positively Modulated by a WYL-Domain-Article Cas13d Is a Compact RNA-Targeting Type VI CRISPR Effector Positively Modulated by a WYL-Domain-Containing Accessory Protein. 327–339 (2018) doi:10.1016/j.molcel.2018.02.028.

16. Acharya, S. et al. Francisella novicida Cas9 interrogates genomic DNA with very high specificity and can be used for mammalian genome editing. Proc. Natl. Acad. Sci. 116, 20959–20968 (2019).

17. Chen, F. et al. Targeted activation of diverse CRISPR-Cas systems for mammalian genome editing via proximal CRISPR targeting. Nat. Commun. 8, (2017).

18. Hirano, H. et al. Structure and Engineering of Francisella novicida Cas9. Cell 164, 950–961 (2016).

19. Chen, J. S. et al. targeting accuracy. Nat. Publ. Gr. 550, 407–410 (2017).

20. Petherick, A. World Report Developing antibody tests for SARS-CoV-2. Lancet 395, 1101–1102 (2019).

21. Babiker, A., Myers, C. W., Hill, C. E. & Guarner, J. SARS-CoV-2 Testing. Am. J. Clin. Pathol. 1–3 (2020) doi:10.1093/ajcp/aqaa052.

22. Guo, Y.-R. et al. The origin, transmission and clinical therapies on coronavirus disease 2019 (COVID-19) outbreak–an update on the status. Mil. Med. Res. 7, 1–10 (2020).

23. Andrieu-Soler, C. & Soler, E. When basic science reaches into rational therapeutic design: from historical to novel leads for the treatment of β-globinopathies. Curr. Opin. Hematol. 27, (2020).

24. Rees, D. C. Sickle Cell Disease. 1561–1573 (2017) doi:10.1056/NEJMra1510865.

25. Landrum, M. J. et al. ClinVar: Public archive of relationships among sequence variation and human phenotype. Nucleic Acids Res. 42, 980–985 (2014).

26. Ribeiro, M. L. et al. Mutations in the 23S rRNA gene are associated with clarithromycin resistance in Helicobacter pylori isolates in Brazil. 4, 1–4 (2003).

27. Binkowska, A., Biernat, M. M., Laczmanski, L. & Gosciniak, G. Molecular Patterns of Resistance Among Helicobacter pylori Strains in South-Western Poland. Front. Microbiol. 9, 1–10 (2018).

28. Perna, S. et al. Association of the bitter taste receptor gene TAS2R38 (polymorphism RS713598) with sensory responsiveness, food preferences, biochemical parameters and body-composition markers. A cross-sectional study in Italy. Int. J. Food Sci. Nutr. 69, 245–252 (2018).

29. Zhao, Y., Chen, F., Li, Q., Wang, L. & Fan, C. Isothermal Amplification of Nucleic Acids. Chem. Rev. 115, 12491–12545 (2015).

30. Suppiah, J. et al. Clinical manifestations of dengue in relation to dengue serotype and genotype in Malaysia: A retrospective observational study. PLoS Negl. Trop. Dis. 12, 1–20 (2018).

31. Vicente, C. R. et al. Serotype influences on dengue severity: A cross-sectional study on 485 confirmed dengue cases in Vitória, Brazil. BMC Infect. Dis. 16, 1–7 (2016).

32. Lobato, I. M. & O’Sullivan, C. K. Recombinase polymerase amplification: Basics, applications and recent advances. TrAC -Trends Anal. Chem. 98, 19–35 (2018).

33. Lewnard, J. A. & Lo, N. C. Comment Scientific and ethical basis for social-distancing interventions against COVID-19. Lancet Infect. Dis. 3099, 2019–2020 (2020).

34. Chan, J. F.-W. et al. Improved molecular diagnosis of COVID-19 by the novel, highly sensitive and specific COVID-19-RdRp/Hel real-time reverse transcription-polymerase chain reaction assay validated in vitro and with clinical specimens. J. Clin. Microbiol. JCM.00310-20 (2020) doi:10.1128/JCM.00310-20.

35. van Doremalen, N. et al. Aerosol and Surface Stability of SARS-CoV-2 as Compared with SARS-CoV-1. N. Engl. J. Med. 0, null.

36. Landrum, M. J. et al. ClinVar: improving access to variant interpretations and supporting evidence. Nucleic Acids Res. 46, D1062–D1067 (2017).

37. Wu, F. et al. A new coronavirus associated with human respiratory disease in China. Nature 579, 265–269 (2020).

38. Snijder, E. J. et al. Unique and Conserved Features of Genome and Proteome of SARS-coronavirus, an Early Split-off From the Coronavirus Group 2 Lineage. J. Mol. Biol. 331, 991–1004 (2003).

39. Brister, J. R., Ako-adjei, D., Bao, Y. & Blinkova, O. NCBI Viral Genomes Resource. Nucleic Acids Res. 43, D571–D577 (2014).

40. Jiang, H. & Wong, W. H. SeqMap: mapping massive amount of oligonucleotides to the genome. Bioinformatics 24, 2395–2396 (2008).

41. Li, H. & Durbin, R. Fast and accurate long-read alignment with Burrows– Wheeler transform. Bioinformatics 26, 589–595 (2010).

42. Quinlan, A. R. & Hall, I. M. BEDTools: a flexible suite of utilities for comparing genomic features. Bioinformatics 26, 841–842 (2010).

