## Supplementary Figure 1 for "Rapid, field-deployable nucleobase detection and identification using FnCas9"

**A**

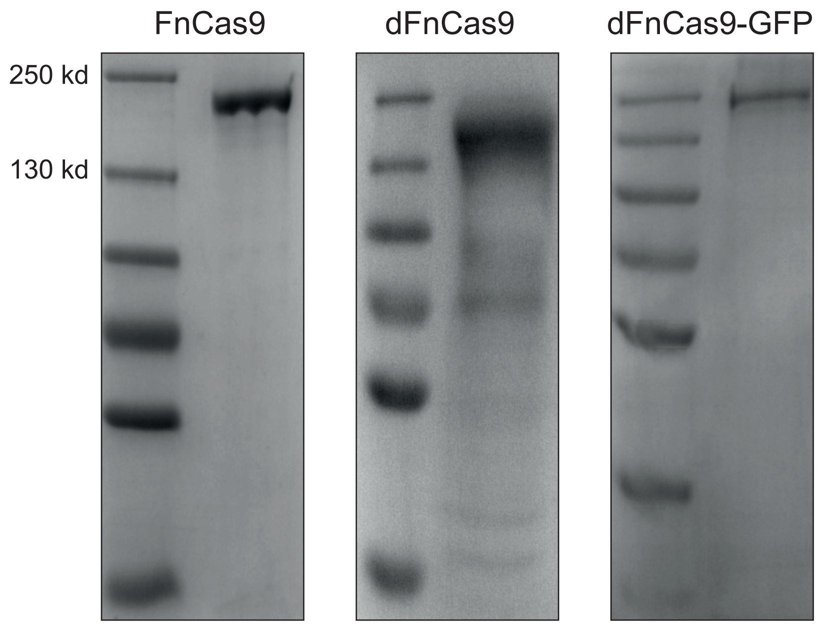


**B**

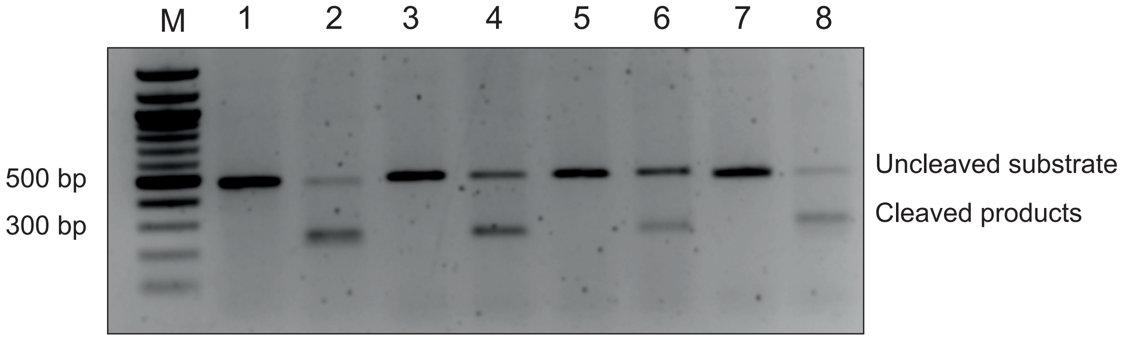


1. WT Glanzmann's Thrombasthenia
2. mut Glanzmann's Thrombasthenia
3. WT Hemophilia A (Factor VIII deficiency)
4. mut Hemophilia A (Factor VIII deficiency)
5. WT Glycogen Storage Disease Type I
6. mut Glycogen Storage Disease Type I
7. WT X- linked myotubular myopathy
8. mut X- linked myotubular myopathy
