## Supplementary Figure 2 for "Rapid, field-deployable nucleobase detection and identification using FnCas9"

Junction for analysis and target design for Your FELUDA assay (JATAYU)

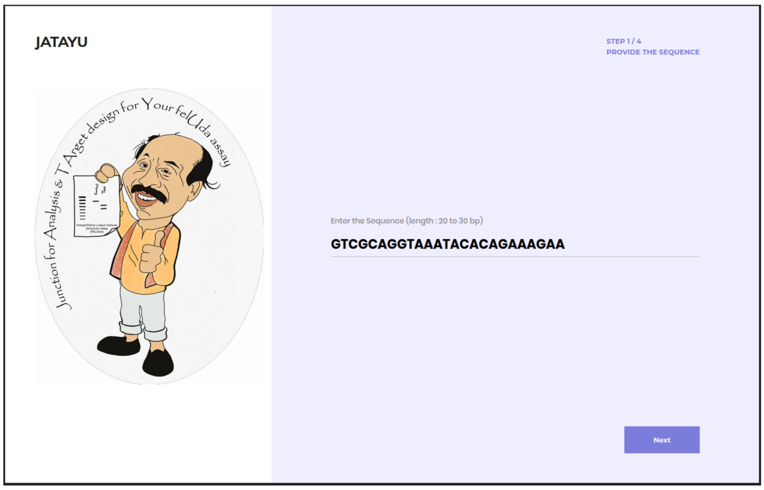


Step 1

Sequence Input

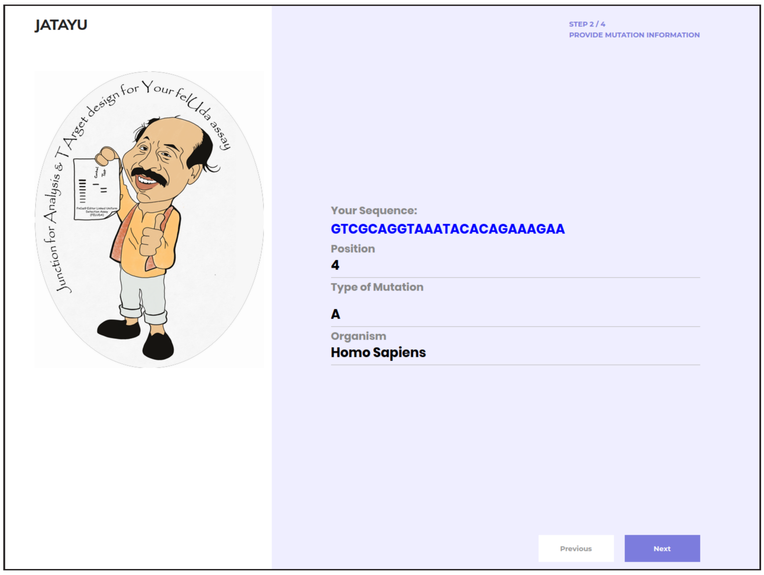


Step 2

Mismatch information

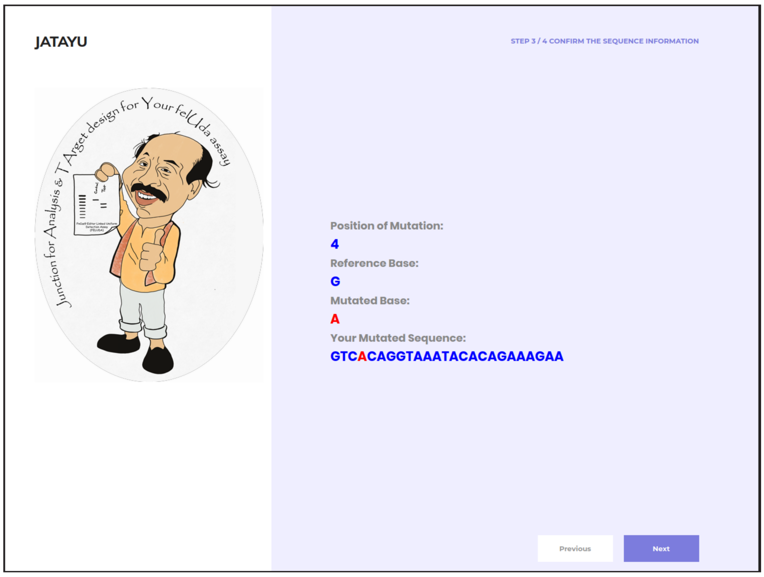


Step 3

Confirmation of data entry

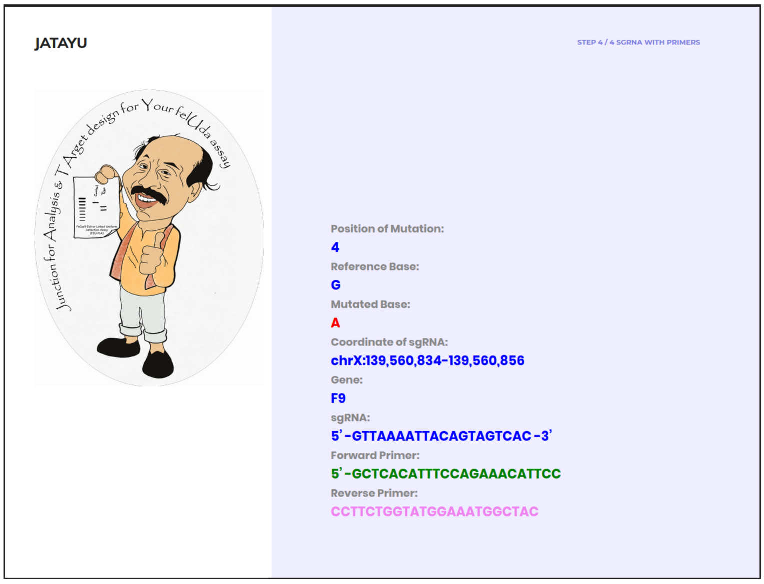


Step 4

Output
