## Supplementary figures and images for "Rapid, field-deployable nucleobase detection and identification using FnCas9"

### Supplementary Figure 3

#### Supplementary Figure 3

A

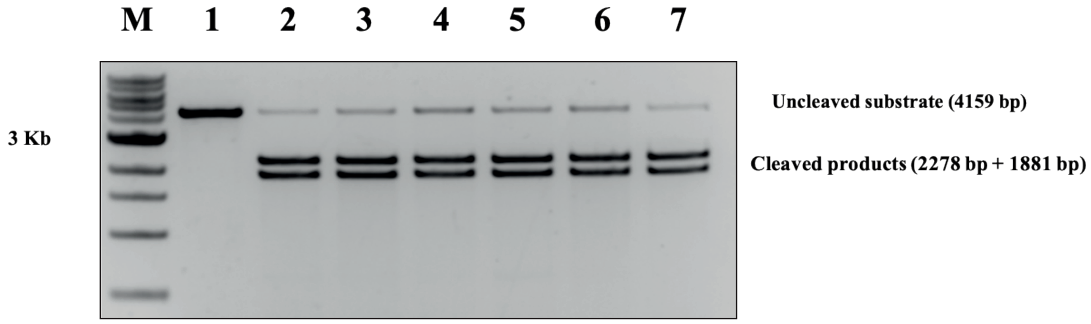


B

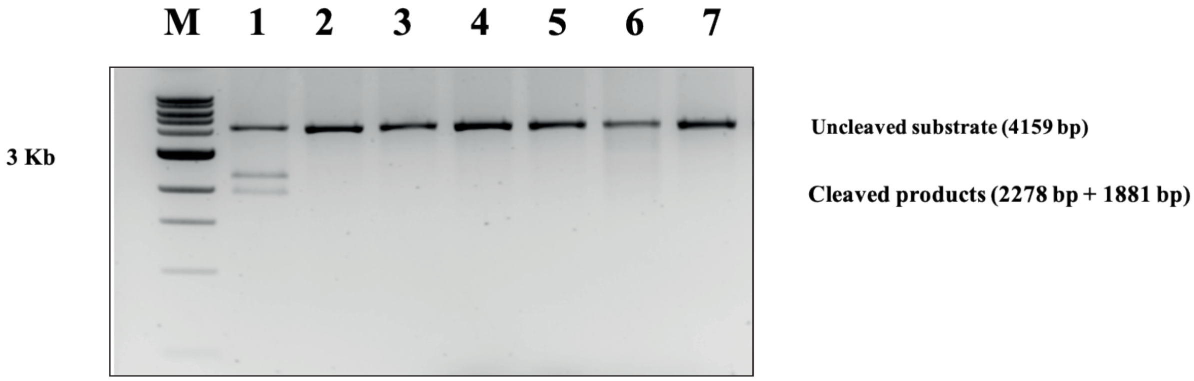

### Supplementary Figure 4

**Supplementary Figure 4**

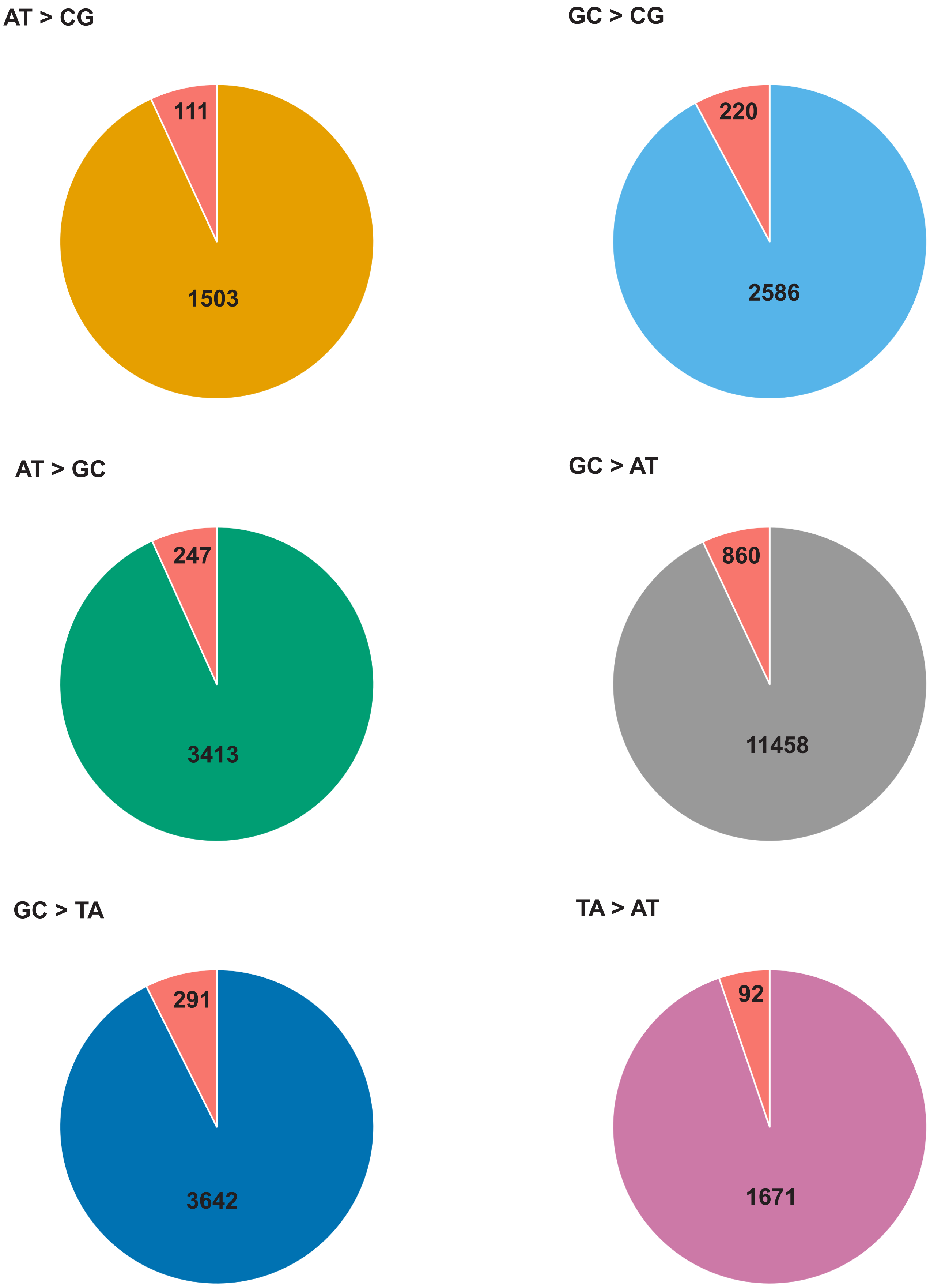

### Supplementary Figure 5

**Supplementary Figure 5**

**A**

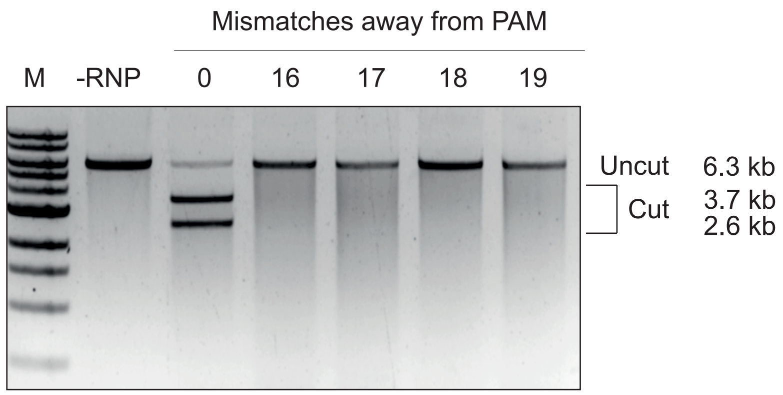


**B**

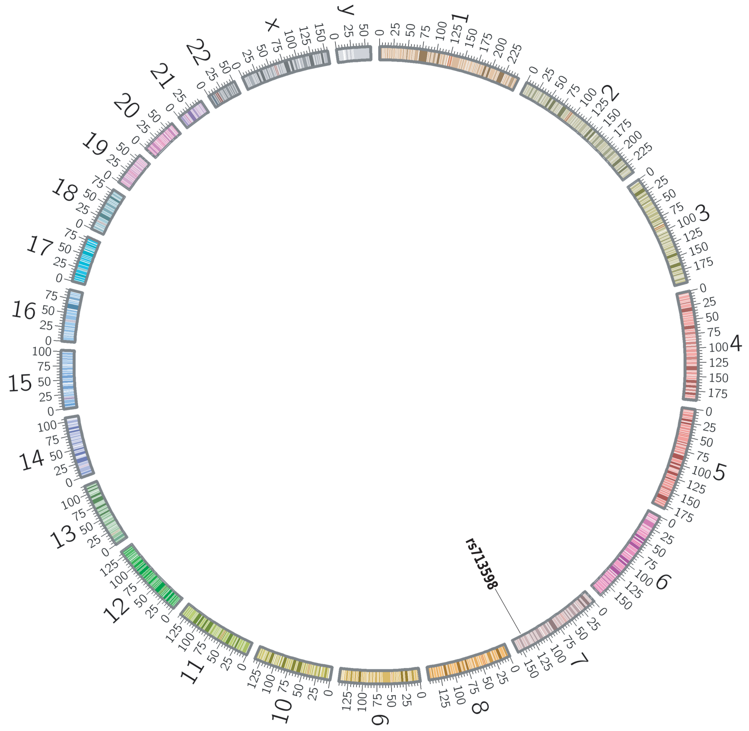


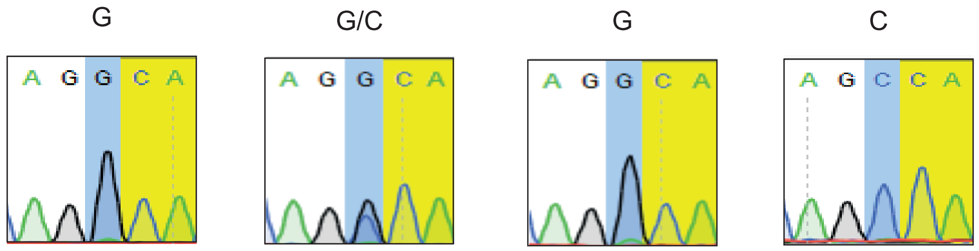


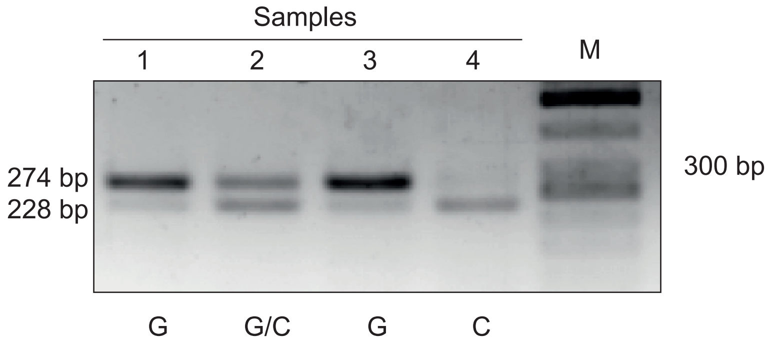

### Supplementary Figure 6

#### Supplementary Figure 6

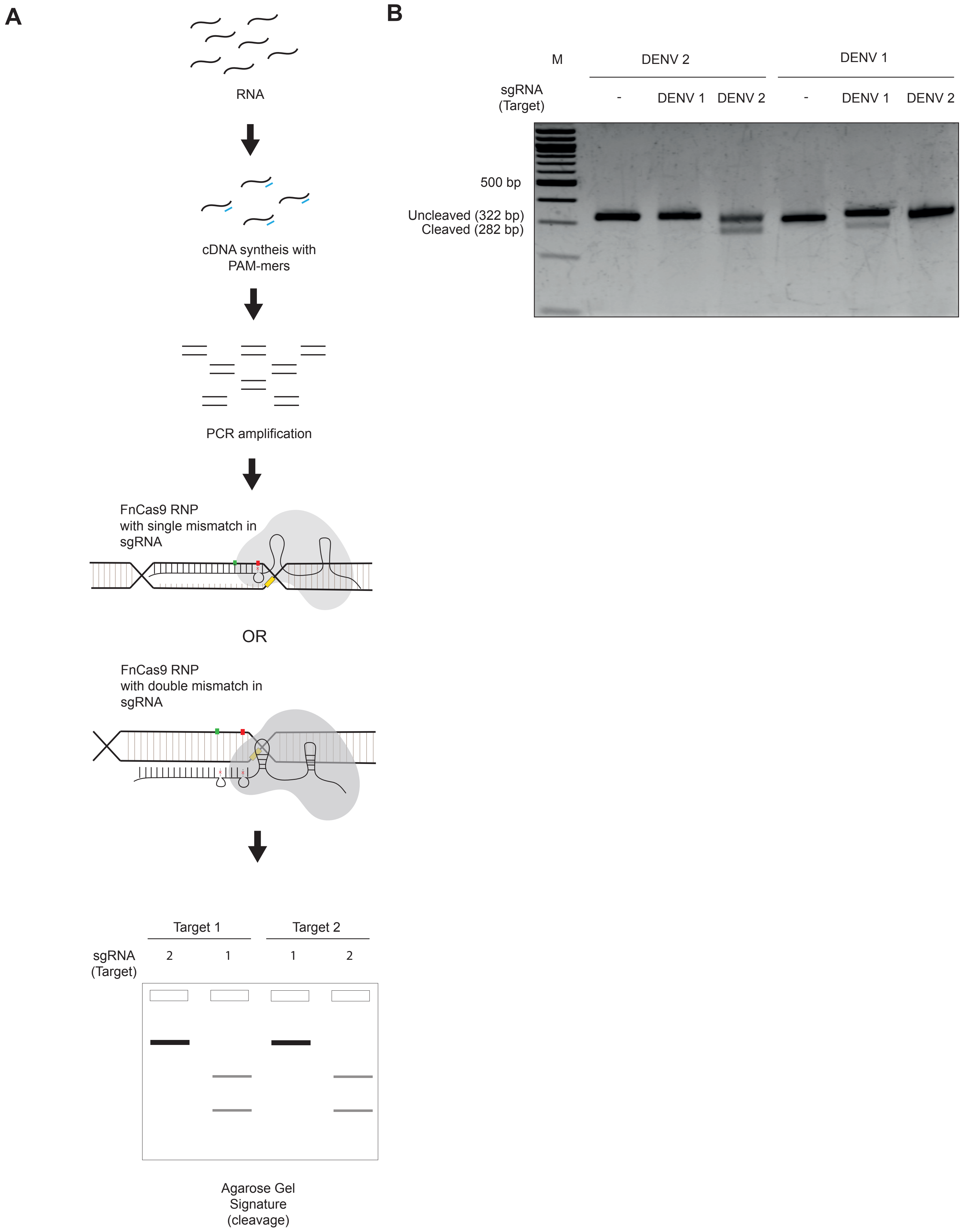

### Supplementary Figure 7

**Supplementary Figure 7**

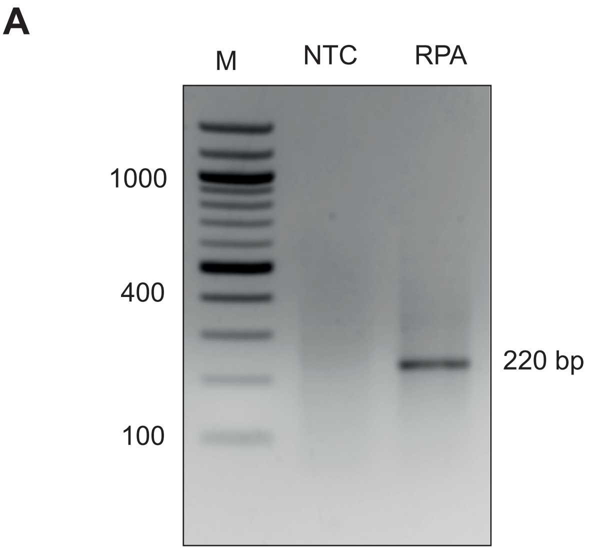


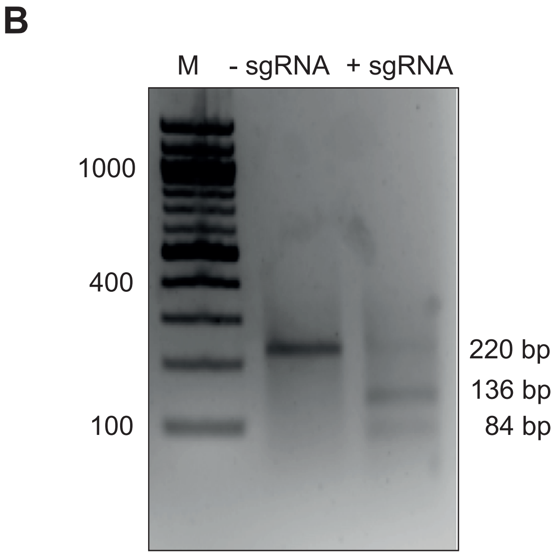

### Supplementary Figure 10

#### Supplementary Figure 10

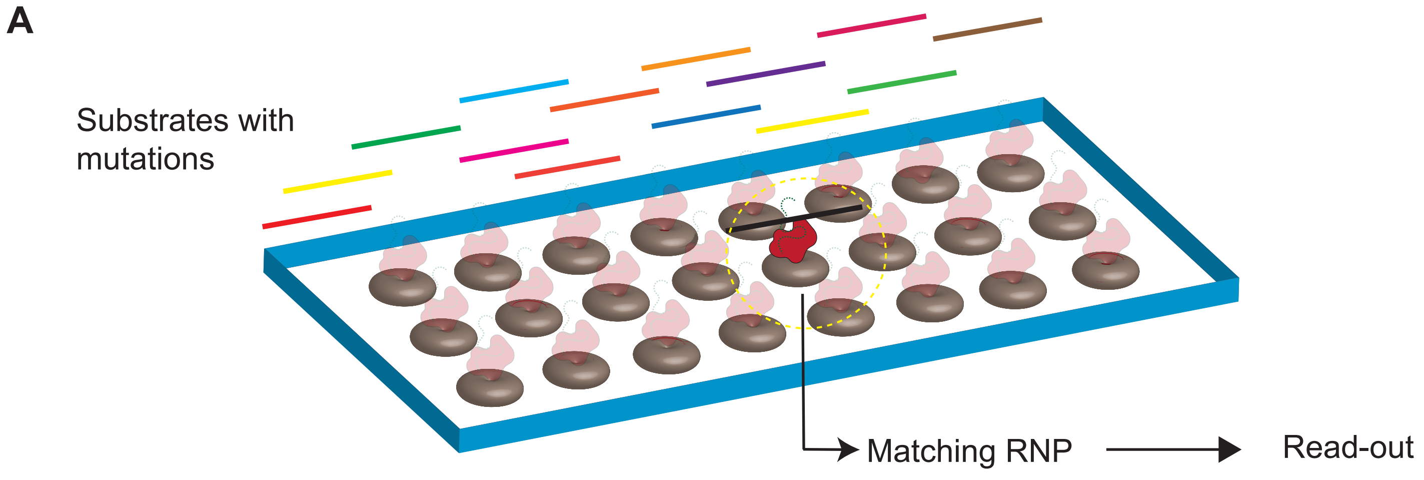


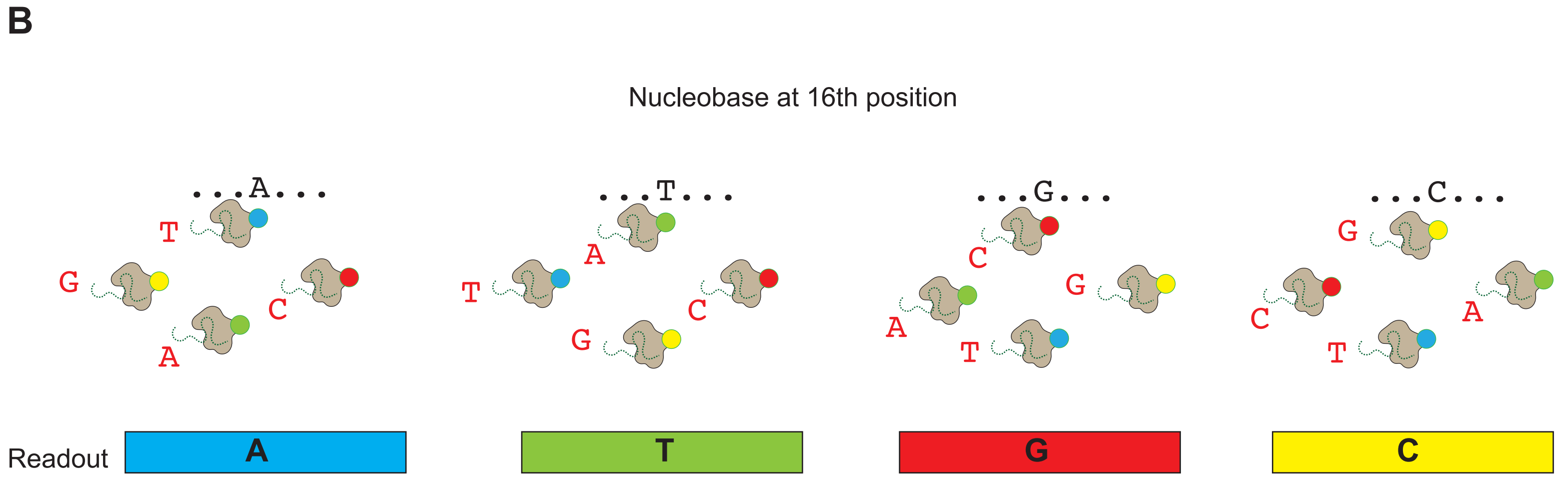
