## Supplementary Figure 8 for "Rapid, field-deployable nucleobase detection and identification using FnCas9"

A

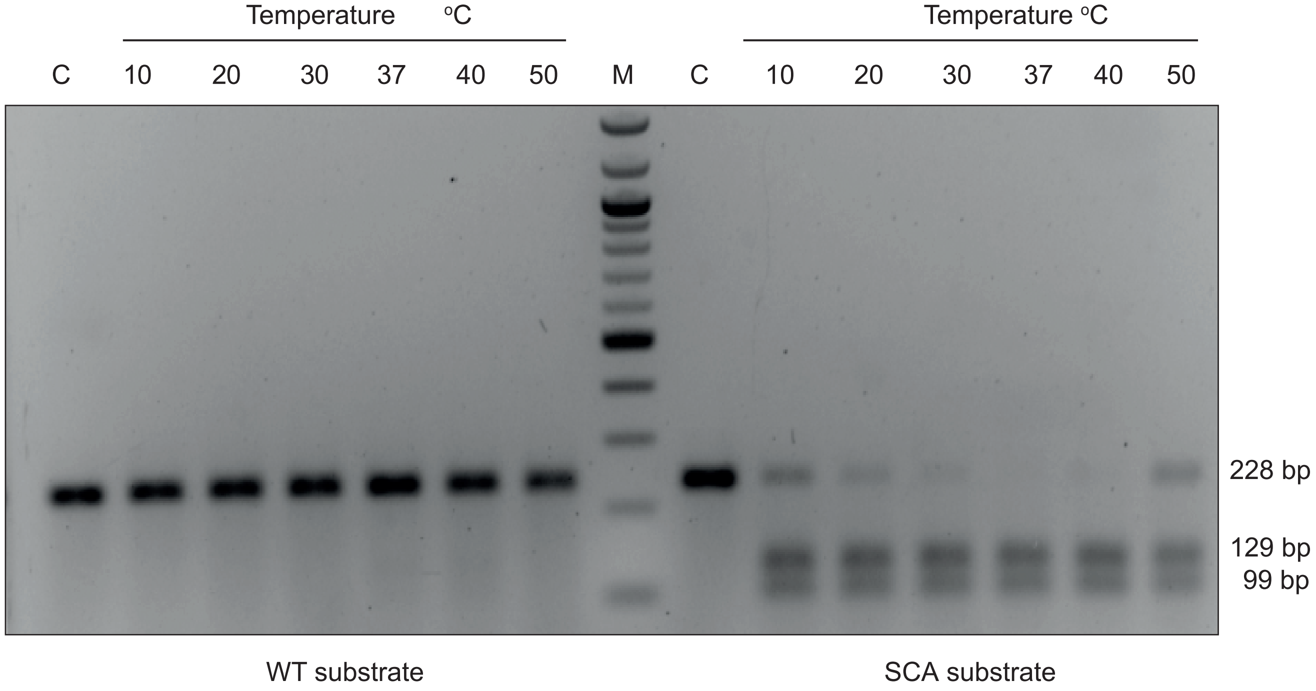


B

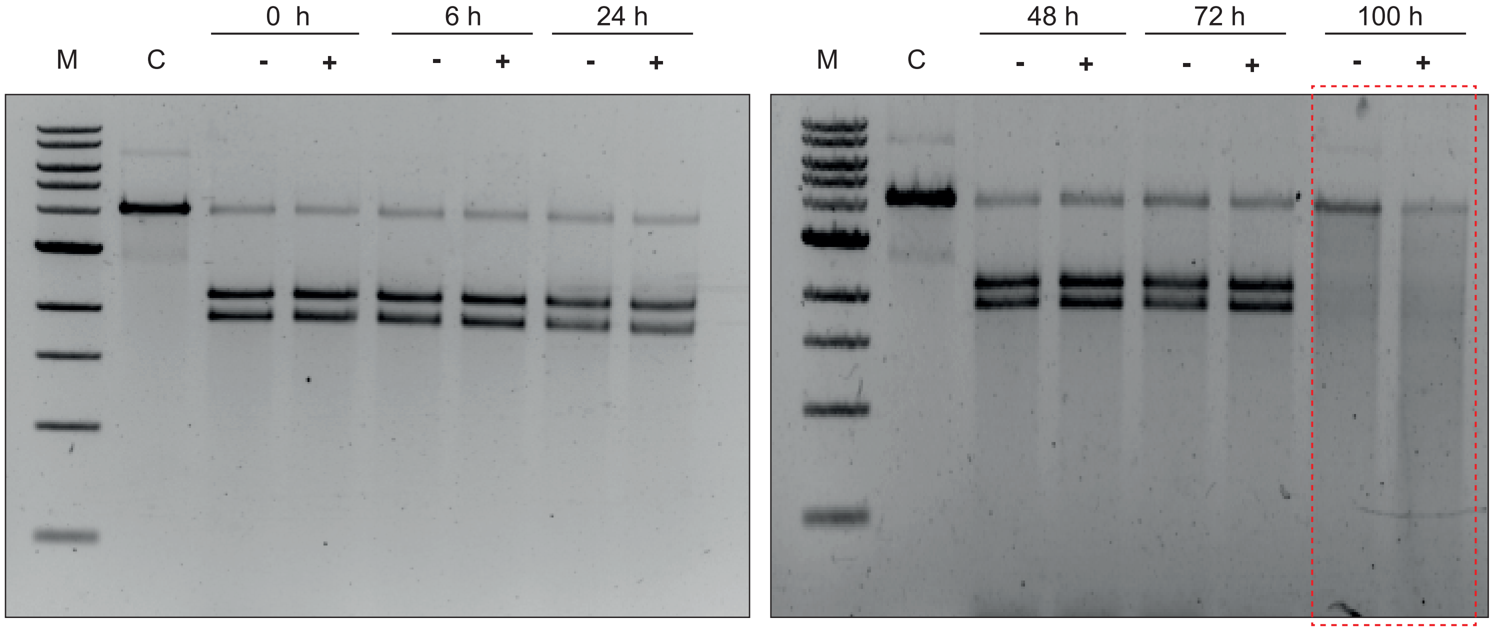


Uncleaved band: 4159 bp

Cleaved bands: 2278 bp
1881 bp

- with 10 % sucrose in Buffer
+ without 10 % sucrose in Buffer
