## Supplementary Figure 9 for "Rapid, field-deployable nucleobase detection and identification using FnCas9"

**A**

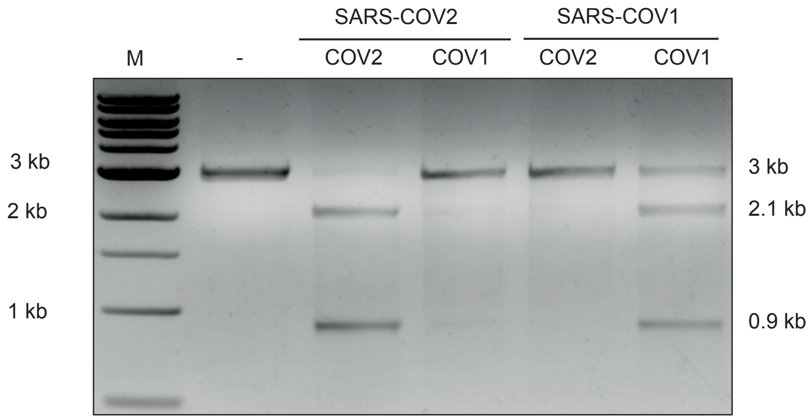


**B**

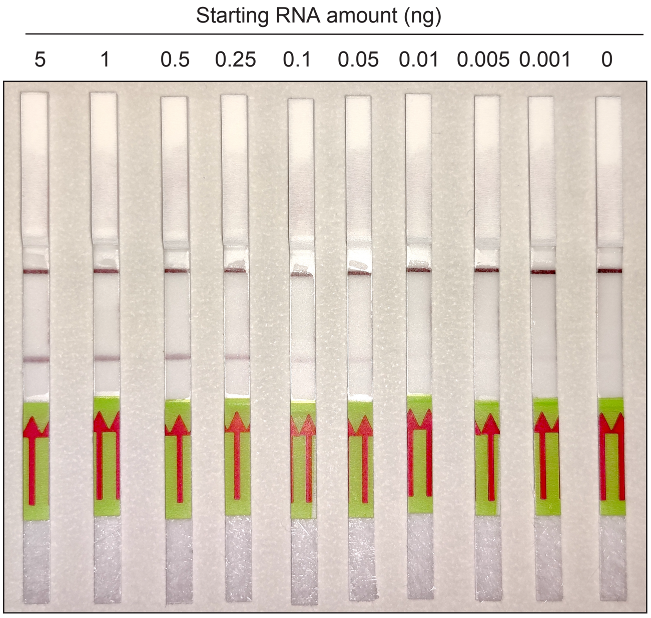


**C**

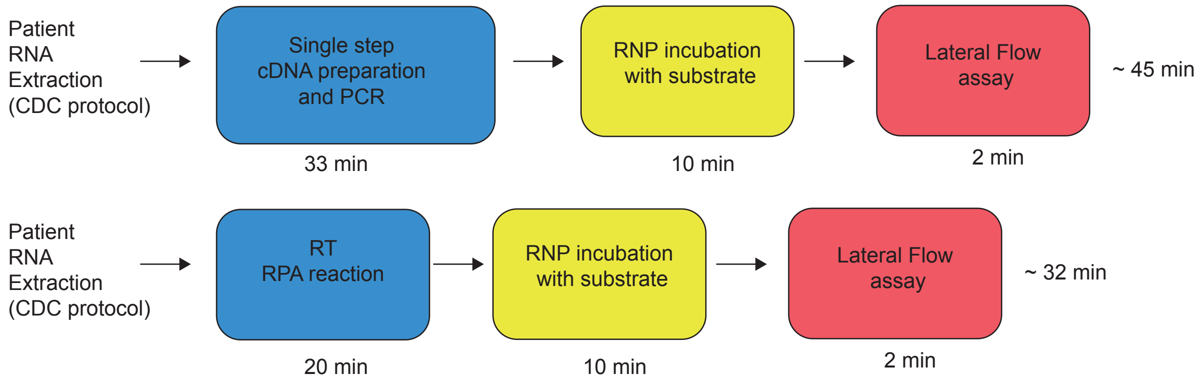


**D**

| Reagents | Cost per reaction (USD) |
|---|---|
| Reverse Transcription reagents | 0.46 |
| DNA Polymerase | 0.20 |
| Biotin labelled primers | 0.01 |
| crRNA synthesis & purification | 0.09 |
| FAM labelled tracrRNA | 0.06 |
| Paper strip | 4.5 |
| FnCas9 protein | 0.03 |
| Total | 5.35 |
