## Supplementary Note 1 for "Rapid, field-deployable nucleobase detection and identification using FnCas9"

**A protocol for rapid detection of the 2019 novel coronavirus (COVID-19) using CRISPR diagnostics: FELUDA**

Preparation of CRISPR-Cas9 crRNAs (to be performed prior to sample handling under RNase free condition):

1. Synthesis of *in vitro* transcribed (IVT) crRNAs targeting COVID-19 and Beta-Actin (ACTB)

- Assemble equimolar ratio of Forward and Reverse oligos (refer Table 1) for each target:

| **Reagents** | **Volume (µl)** | **Final Concentration** |
|---|---|---|
| Forward Oligo (100 µM) | 1.25 | 2.5 µM |
| Reverse Oligo (100 µM) | 1.25 | 2.5 µM |
| Total Volume (with nuclease free water) | upto 50 µl | |

- Heat the reaction mix at 95ºC for 5 min followed by slow cooling at room temperature for 15 min.
- Perform *in vitro* transcription using commercially available T7 Polymerase based IVT kit as per recommended protocol. A sample is given below (MEGAscript T7 Kit, ThermoFisher Scientific)

  Assemble the following reaction components at room temperature:

| **Reagents** | **Volume (µl)** | **Final Concentration** |
|---|---|---|
| Nuclease free water | upto 20 | |
| ATP (75 mM) | 2 | 7.5 mM |
| GTP (75 mM) | 2 | 7.5 mM |

| | | |
|---|---|---|
| CTP (75 mM) | 2 | 7.5 mM |
| UTP (75 mM) | 2 | 7.5 mM |
| 10X Reaction Buffer | 2 | 1X |
| Enzyme Mix | 2 | - |
| Annealed oligo duplex from step 1 | 5 | - |

- Incubate the mix overnight at 37°C.
- Add 1 µl of Turbo DNAse in the reaction mix and incubate at 37°C for 30 min.
- Heat inactivate at 70°C for 10 min.

  **Optional**: RNA can be visualized on gel (2% agarose) to check for its integrity.

  Column based RNA clean-up (such as NucAway Spin Columns, AM10070, ThermoFisher Scientific)

2. Generation of chimeric gRNAs (crRNA:tracrRNA-FAM)

| Reagents | Volume (µl) | Final Concentration |
|---|---|---|
| IVT synthesized crRNA | - | 2 µM |
| FAM labelled tracrRNA | - | 2 µM |
| Annealing Buffer (100 mM NaCl, 50 mM Tris-Cl pH 8.0, 1 mM $MgCl_2$) | upto 50 | |

- Heat the reaction mix at 95°C for 5 min followed by slow cooling at room temperature for 15 min.

The crRNA and chimeric gRNA products can be produced in bulk and stored at -20°C for long term use.

**FELUDA based detection of COVID-19 from patient samples:**

1. Extract patient RNA according to CDC recommendations.

2. Set-up single step RT and PCR reaction.

Assemble the reaction components as below (using Biotin labelled primers):

| **Reagents** | **Volume (µl)** | **Final Concentration** |
|---|---|---|
| Forward Biotinylated Primer (10 µM) | 0.2 | 200 nM |
| Reverse Biotinylated Primer (10 µM) | 0.2 | 200 nM |
| dNTPs (2.5 mM) | 0.4 | 100 µM |
| 10X Reaction Buffer (200mM Tris-Cl pH 8.4, 500mM KCl) | 1 | (1X) |
| $MgCl_2$ (50 mM) | 0.3 | 150 µM |
| Taq DNA polymerase (5U/µl) | 0.02 | 0.01 U/µl |
| Reverse Transcriptase (200U/µl) | 0.2 | 4 U/µl |
| RNA sample (0.5-5ng) | As per sample | |
| RNase inhibitor (Optional, in case not present with RT enzyme) 20U/µl | 0.2 | 0.4 U/µl |
| Total Volume (with nuclease free water) | upto 10 | |

Tip: During optimization, an aliquot of the PCR product may be run on gel to check successful RT-PCR reaction conditions.

**Reaction Conditions:**

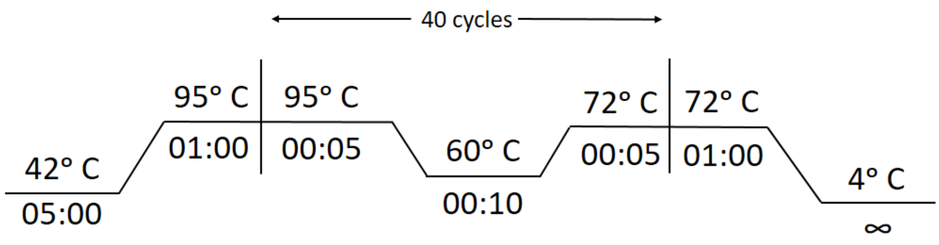


3. Prepare dFnCas9-chimeric gRNA-RNP complexes for the samples to be tested. One complex each for COVID-19 gene (positive) and ACTB (negative) may be assembled for each sample.

- Incubate dFnCas9 with Chimeric FAM labelled guide RNA to generate RNP complexes for 10 min at Room Temperature (RT).

| **Reagents** | **Volume (µl)** | **Final Concentration** |
|---|---|---|
| dFnCas9 (1µM) | 0.5 | 50 nM |
| Chimeric FAM labelled gRNA (2 µM) | 0.25 | 50 nM |
| Total Volume (with Buffer containing 20 mM HEPES pH 7.5, 150 mM KCl, 10% glycerol, 1mM DTT and 10mM $MgCl_2$) | upto 9 | |

4. Combine 1 µL of the biotinylated amplicon (from Step 2) with 9µL of the dFnCas9 RNP complexes (from Step 3)

5. Incubate the complex at 37°C for 15 minutes in a heating block or water bath.

6. Insert Milenia HybriDetect 1 lateral flow strip directly into reaction tubes **(for positive, COVID-19 CRISPR RNP and negative ACTB CRISPR RNP complexes)**

7. Allow the solution to migrate into the strip for 2 minutes at room temperature and observe the result.

- **Beta-Actin (ACTB) crRNA to be used as negative control.**
- **For accurate results, positive samples show up in the test band within 2 min while negative samples show very weak or no signal.**
- Incubating for longer times leads to increasing background signal intensity at test band location making interpretation ambiguous.
- Signals between positive and negative assays can also be interpreted by densitometry analysis from images captured using any photographic device.

8. Strips may be discarded according to standard procedures.

**Table 1: List of primers used**

| Primer | Sequence (5'-3') |
|---|---|
| COVID_cr1_FP-Biotin | Biotin-CAGGCTGTTGCTAATGGTGA |
| COVID_cr1_RP Biotin | Biotin-TGTTCAAGGGAACACAACCA |
| COVID_cr2_FP-Biotin | Biotin-TTGCCAGGAACCTAAATTGG |
| COVID_cr2_RP-Biotin | Biotin- GAATCTGAGGGTCCACCAAA |
| ACTB_crRNA1_Forward_oligo | TAATACGACTCACTATAGCCGCGCTCGTCGTCGACAAGTTTCAGTTGCTGAATTAT |
| ACTB_crRNA1_Reverse_oligo | ATAATTCAGCAACTGAAACTTGTCGACGACGAGCGCGGCTATAGTGAGTCGTATTA |
| ACTB_crRNA2_Forward_oligo | TAATACGACTCACTATAGGATAGCAACGTACATGGCTGTTTCAGTTGCTGAATTAT |
| ACTB_crRNA2_Reverse_oligo | ATAATTCAGCAACTGAAACAGCCATGTACGTTGCTATCCTATAGTGAGTCGTATTA |
| COVID_crRNA1_Forward_oligo | TAATACGACTCACTATAGTATAAACAGGCTAGATCTGGTTTCAGTTGCTGAATTAT |

| | |
|---|---|
| COVID_crRNA1_Reverse_oligo | ATAATTCAGCAACTGAAACCAGATCTAGCCTGTTTATACTATAGTGAGTCGTATTA |
| COVID_crRNA2_Forward_oligo | TAATACGACTCACTATAGGTCCACCAAACGTAATGCGGTTTCAGTTGCTGAATTAT |
| COVID_crRNA2_Reverse_oligo | ATAATTCAGCAACTGAAACCGCATTACGTTTGGTGGACCTATAGTGAGTCGTATTA |
| TracrRNA-FAM | G*U*AAUUAAUGCUCUGUAAUCAUUUAAAAGUAUUUUGAACGGACCUCUGUUUGACACGUC*U*G-FAM |

* Phosphorothioate bond

**In vitro transcription reagents used:**

MEGAscript T7 Transcription Kit (AM1334, ThermoFisher Scientific)

**RT Enzymes that have been successfully tested:**

1. RevertAid RT (Cat. EP0441, ThermoFisher Scientific)
2. Quantitect RT (Cat. 205311, Qiagen)
3. iScript RT (Cat. 1708890, BIO-RAD)
4. High Capacity cDNA RT (Cat.  4368814, ThermoFisher Scientific)
5. Reverse Transcriptase Core Kit (Cat. RT-RTCK-03, Eurogentec)

**DNA Polymerase used:**

1. Taq DNA Polymerase (Cat. 18038018, ThermoFisher Scientific)
2. Taq DNA Polymerase (Cat. MB101-0500, GeneDireX, Inc.)
3. Taq DNA Polymerase (Cat. # TQ050, Geneaid Biotech)
4. DyNAzyme II DNAPolymerase (Cat. #F-501L, ThermoFisher Scientific)
5. KAPA Taq PCR Kit (Cat. KK1015, Sigma-Aldrich)

**Biotinylated primers:**

Commercial vendors such as BioServe Biotechnologies (Hyderabad, India) or Merck (Darmstadt, Germany) can provide custom labelled oligos.

**Synthetic labelled tracRNA:**

6 – Fluorescein amidite (6-FAM) labelling at 3' end of FnCas9 tracrRNA can be commercially purchased from GenScript Biotech (New Jersey, USA) or Merck (Darmstadt, Germany). This oligo is 61 bases in length.

**Paper strip for detection:**

MILENIA HybriDetect1 (Cat. No. MILENIA01, TwistDx, UK or Giessen, Germany).

**Single step RT and PCR have been successfully tested with following Thermal Cycler:**

1. Veriti 96-Well Thermal Cycler (Cat. No. 4375786, ThermoFisher Scientific).

2. ProFlex 3 x 32-well PCR System (Cat. No. 4484073, ThermoFisher Scientific).

3. Applied Biosystems 2720 Thermal Cycler (Cat. No. 4359659, ThermoFisher Scientific).

3. T100 Thermal Cycler (Cat. No. 186-1096, BIO-RAD).
